# Comparison of missing data handling methods for variant pathogenicity predictors

**DOI:** 10.1101/2022.06.17.496578

**Authors:** Mikko Särkkä, Sami Myöhänen, Kaloyan Marinov, Inka Saarinen, Leo Lahti, Vittorio Fortino, Jussi Paananen

## Abstract

1

**Background:** Modern clinical genetic tests utilize next-generation sequencing (NGS) approaches to comprehensively analyze genetic variants from patients. Out of these millions of variants, clinically relevant variants that match the patient’s phenotype need to be identified accurately within a rapid timeframe that facilitates clinical action. As manual evaluation of variants is not a feasible option for meeting the speed and volume requirements of clinical genetic testing, automated solutions are needed. Various machine learning (ML), artificial intelligence (AI), and *in silico* variant pathogenicity predictors have been developed to solve this challenge. These solutions rely on the comprehensiveness of the available data and struggle with the sparse nature of genetic variant data. Therefore, careful treatment of missing data is necessary, and the selected methods may have a huge impact on the accuracy, reliability, speed and associated computational costs.

**Results:** We present an open-source framework called AMISS that can be used to evaluate performance of different methods for handling missing genetic variant data in the context of variant pathogenicity prediction. Using AMISS, we evaluated 14 methods for handling missing values. The performance of these methods varied substantially in terms of precision, computational costs, and other attributes. Overall, simpler imputation methods and specifically mean imputation performed best.

**Conclusions:** Selection of the missing data handling method is crucial for AI/ML-based classification of genetic variants. We show that utilizing sophisticated imputation methods is not worth the cost when used in the context of genetic variant pathogenicity classification.

## 2 Introduction

### 2.1 Genetic variant pathogenicity prediction

Adoption of next-generation sequencing (NGS) technology has greatly improved the scalability of genetic sequencing in both research and clinical genetics. Whole-exome and whole-genome sequencing (WES and WGS) are now becoming standard methodology, allowing detection of variants in a much broader set of loci than was previously feasible. However, the large numbers of detected variants from each sample present problems especially in clinical contexts. The often time-critical process of identifying clinically relevant genetic variants requires the manual and painstaking exploration of long variant lists. Highly accurate computational tools could further improve the filtering or prioritization process.

Tools applicable for this purpose already exist. Many of them were developed to be used in research contexts, though many are also used in clinical variant interpretation (see ACMG/AMP guidelines [1]). *In-silico* variant effect and pathogenicity prediction tools have been developed to inform the selection of variants for further testing (e.g. SIFT [2], PROVEAN [3], MutationTaster2 [4, 5], LRT [6] and FATHMM [7]). Similarly, gene and variant prioritization tools have been developed to rank variants for exploration (e.g. VAAST [8], PHEVOR [9], FunSeq [10], PHIVE [11] and Phen-Gen [12]). Peterson & al. [13], Niroula & Vihinen [14], and Eilbeck & al. [15] discuss the already diverse set of prediction tools.

Peterson & al. note that the primary feature of most deleteriousness prediction tools is a metric of sequence conservation, often supplemented with additional features, even ones derived from outputs of previously developed tools [13]. The inclusion of outputs of several previously developed tools as input to a machine learning (ML) method forms the class of *metapredictors* or *ensemble predictors*.

Examples of metapredictors are REVEL [16], CADD [17, 18], DANN [19], Eigen [20], PON-P [21] and MetaSVM and MetaLR [22]. Metapredictors often report high performance [23, 24]. Metapredictors are often also able to predict outcomes for wide ranges of variants, since they can simultaneously utilize tools that are aimed at variants with different predicted consequences by e.g. combining missense variant effect prediction tools and splicing variant effect tools.

Since traditional tools have been developed using different data, different methods and at various times, they may differ in the set of variants for which prediction is possible, even if their intended use cases match. If one wants to build a metapredictor that incorporates several existing tools but does not want to restrict the domain of prediction, this means that the input features for metapredictors have *missing values*. Missing values can also arise in features representing experimentally obtained data. For example, when incorporating allele frequency information from gnomAD [25], variants that were not observed in the aggregated cohorts will have a missing value as their allele frequency.

### 2.2 Aim

The high prevalence of missing values in annotated variant data implies that the chosen method for handling missingness will have a major impact on the performance of variant pathogenicity metapredictors and ML/AI based variant classification and interpretation tools. Our aim is to identify missingness handling methods most likely to produce good results in this context. This includes comparing achieved classification performance, difficulty of implementation, and computation time. For this purpose, we present a framework implemented in the `R` language [26] that treats variant data with a variety of missingness handling methods and then trains and evaluates a ML classifier on each treated dataset.

We focus on the comparison of imputation methods, as they are the most generally applicable and allow use of any ML/AI method on the imputed dataset. These properties are shared by the missingness indicator augmentation method, which is also included. We limit our evaluation to imputation of numerical features, as categorical features have a relatively natural treatment through an additional category denoting missingness.

We perform three experiments based on executing the above-mentioned framework on the ClinGen [27] subset of variants from ClinVar [28, 29, 30]. In the first experiment, we repeatedly simulate additional missing values in the training dataset. We then use these generated datasets and the framework to evaluate, for each imputation method,

1. the effect of increasing missingness on classification performance,
2. the relationship between imputation error, which is the difference between original known values and imputed values, and classification performance.

In the second experiment, we perform repeated random sub-sampling crossvalidation on the training set to assess whether there are differences in the methods’ susceptibility to changes in dataset composition.

In the third, main experiment, we use the full data and wider parameter grids to execute the framework to obtain a realistic estimate of the effect of imputation method choice in variant pathogenicity metapredictor construction.

## 3 Background

### 3.1 Missing data

Data with and without missing values are often called *incomplete* and *complete* data, respectively. Consider a matrix of *A* that represents the unobserved, underlying values that would be obtained by data collection in the absence of any missing data generation mechanisms. The subset of values of *A* that are observed in data collection is denoted *A*_*obs*_, and the subset of missing values of *A* is denoted *A*_*mis*_. The values of *A*_*mis*_ are not known when analysing any real dataset. *M* is the missingness indicator matrix whose values are 0 when the corresponding value of *A* is observed, and 1 when the corresponding value of *A* is missing.

Traditionally, missing data mechanisms are classified into *missing completely at random* (MCAR), *missing at random* (MAR) and *missing not at random* (MNAR) [31]. In a missing data process with an MCAR mechanism, the probability of a value being missing does not depend on any observed or unobserved values. With an MAR mechanism, the probability of a value being missing may depend on observed values, and with an MNAR mechanism, the probability of a value being missing may depend on both observed and unobserved values. Van Buuren notes data from a truly random sample of a population as an example of MCAR [32, subchapter 1.2]; measurements for individuals that were not selected in the sample do not depend on the underlying values of the individual or the observed values of other individuals. Practical examples of MCAR are also incidents unrelated to biology, such as freezer or hard disk failures or file corruption. For non-MCAR missingness gnomAD [25] allele frequency can be used as an example. When annotating samples with allele frequencies, a missing value may indicate that the variant was not present in the study population used to compute the allele frequencies, and thus imply that it might also be extremely rare in healthy individuals. Very low allele frequencies thus end up with missing values and exhibit MNAR behavior. Alternatively, it might indicate the variant is present but was not observed because it resided in a difficult-to-sequence locus and thus exhibited MAR behavior dependent on position.

The above classification is essential in statistical inference, and the validities of different methods depend on which mechanisms appear in the data. However, Sarle [33] notes that “The usual characterizations of missing values as *missing at random* or *missing completely at random* are important for estimation but not prediction”, and Ding & Simonoff [34] provide evidence in support of this statement in the use of classification trees. In an interesting reversal, the presence of *informative missingness* [33, 34] *in the data (i*.*e. missingness being dependent on the response variable conditional on A*_*obs*_) may lead to improved predictive accuracy compared to complete data [34].

#### 3.1.1 Missingness handling in prediction

Rather than inferring properties of the statistical distribution of variant annotations, our aim is production of predictions of pathogenicity, be it a classification into pathogenic and non-pathogenic, a class probability or some other metric. Our aim is thus not statistical inference, for which most missingness handling methods have been developed. It is not clear how the performance of a missingness handling method in one context relates to its performance in another. Overall, there is a significant distinction between these aims, which leads to a variety of differences in the concerns that one must account for [35, 36].

Most studies introducing imputation methods focus on problems where missingness may occur in the model estimation phase, but where data is assumed to be complete at prediction time. Similarly, comparisons of missingness handling methods in machine learning tend to consider missing values only in the context of the training set. In this context, it suffices to design methods which facilitate model training in a way that maximizes generalization performance. In our use case, missing values abound also in the test set, and as such any methods must also allow treatment of the data on which we predict.

This raises additional challenges. Most imputation methods are not implemented in a way that easily allows reuse of learned parameters. That is, it is difficult to first estimate imputation method parameters on the training set, and then impute the test set using the same parameters. This leads to diminished prediction accuracy, as the distributions of imputed data differ in the training and test sets. Even worse, since the parameters for test set imputation are estimated from the test set, the content and size of the test set itself may affect the predictions for individual observations and make it impossible to predict one observation at time.

#### 3.1.2 Methods for missingness handling

The majority of both machine learning models and statistical models are unable to directly use data with missing values. To use such data, one can adopt one of the following strategies.

##### Removal of incomplete observations

Simple removal of incomplete observations is a valid approach when all data we wish to predict on can be assumed to be complete. This is not a viable alternative for variant pathogenicity prediction, since the predictor would be unable to predict on any variant whose feature vector contains any missing values. The number of such variants grows as more features with missing values are added, and as such would greatly restrict the available set of features.

##### Model-based approach

There exist classification methods that can be trained and can predict directly on feature vectors with missing values. Such models may require explicit specification of the mechanism behind missing values. For example, Marlin [37] presented a variation on linear discriminant analysis (LDA)that can both be trained and predict on feature vectors with missing values. In the class of non-linear models, random forest [38] offers a method for handling missing data by iteratively making use of the proximities of observations [39], but is only available for the training phase. CART offers a missingness handling method [40] via so-called surrogate splits, but this method is not suitable for an ensemble of trees like random forest (see [41]). The `randomForestSRC` R packageimplements a missingness handling strategy that is usable in both training and prediction phases [41].

##### Imputation

Imputation is the process of replacing the missing values in data by explicit values. When done exactly once, after which the imputed (or *completed*) dataset is analyzed in the ordinary fashion, the process is termed *single imputation*. Common single imputation strategies are [31]:

- constant imputation; for example, imputation with zeroes
- unconditional mean imputation, where missing values within the same feature are replaced by the mean of the observed values of that feature
- conditional mean imputation, where replacing values can be means dependent on the observed values of other features; for example, by modeling with regression
- drawing from a predictive distribution, where values can be estimated by, for example, addition of a random deviate to a conditional mean estimate, or by drawing the value randomly from the set of observed values

If aiming to perform statistical inference on the imputed data, one must carefully ensure that, in addition to unbiasedness of any parameter estimates, uncertainty introduced by missingness is correctly reflected in the estimated standard errors. This contrasts with designing methods simply for accurate prediction of the underlying value (which is yet again distinct from only facilitating the accurate prediction of the response variable).

Indeed, naïvely imputing data with a single imputation method is misleading as use of even highly accurate single imputation methods will cause underestimation of the standard errors in inference [32, subchapter 2.6]. The uncertainty can be properly incorporated via two main avenues: *likelihood-based approaches* and *multiple imputation* (see [31]). Likelihood-based approaches estimate the parameter of an assumed probability distribution representing the missingness generating process, and do not require explicit imputation of missing values [31].

As opposed to single imputation, the main idea of multiple imputation for statistical inference is to impute the incomplete data multiple times with draws from predictive distributions, fit separate models on each imputed dataset, and *pool* the parameter estimates [31]. In classification, there are several ways to utilize the set of imputed datasets. The first is to obtain estimates of the variability of the classification performance due to the randomness from draws from a predictive distribution. The second would be that one can in principle train a classifier on each, averaging results in the prediction phase [37, 43]. The benefits of this latter approach are not explored in this paper.

##### Reduced models

A somewhat brute-force approach to missingness handling in classification is to train a separate predictor for each combination of missing features using only the available features for each subset. This approach is called the reduced models [37] or reduced-feature models [44] approach, and is observed by Saar-Tsechansky & Provost to consistently perform well [44]. However, a naïve implementation of reduced models easily leads to extremely high computational time and storage requirements, and the hybrid and on-demand approaches described by Saar-Tsechansky and Provost [44] are not trivial to implement.

##### Missingness indicator augmentation

A conceptually simple method is to add, for each original feature, an extra indicator feature that takes the value 1 in observations with the original feature missing, and 0 when the original feature is observed [45]. The missing values of the original feature are then filled with zeroes, and identical indicator features may be removed. The downside with this method is that it may lead to a significant increase in dimensionality of the data (doubling it in the worst case).

#### 3.1.3 Missingness handling in existing tools

##### Missingness causes and handling in traditional tools

Most missingness in variant annotation data can be attributed to the following causes:

1. Insufficient information related to a sequence
2. Inapplicability to the variant

The first cause describes situations where, for example, protein function change prediction tools based on metrics computed from multiple alignment are unable to find sufficiently many matches to the input sequence. In these cases, the tool has no information on which to base its estimates. Another example of this cause is values estimated from scientific studies (e.g. allele frequencies, functional properties of proteins or expression levels of transcripts). Variants that are unobserved in the cohorts, excluded from genotyping microarrays, or inside hard-to-sequence loci will have no observed allele frequency.

The second cause refers to attempts to annotate a variant with a tool that is inapplicable to its predicted molecular consequence. For example, a tool predicting change in protein function cannot produce an estimate for an intergenic variant.

##### Missingness handling in existing metapredictors

Strategies for missingness handling vary widely between different existing metapredictors. REVEL[16] uses k-NN imputation when a variant’s missingness is *≤*50%, and mean imputation when missingness is *>* 50%. CADD [17, 18] and DANN [19] use a mix of manually defined default values to replace missing values, with added missingness indicators for certain features, and mean imputation for genome-wide measures (see Supplementary information for [17]). M-CAP replaces missing values with constants representing the maximally pathogenic prediction for each component tool [46]. Eigen [20] utilizes separate strategies for training and test phases, and builds several weighted linear combinations of its features depending on variant type. This simplifies the situation by only requiring annotations applicable to a specific variant to be available. Learning the weights is based on pairwise correlation, which can be estimated in the presence of some missing values. In the test phase, Eigen performs mean imputation for features that are applicable to the specific variant. KGGSeq ignores any variants that has missing values in its features [47]. PRVCS [48] removes variants with missing values in the training phase and replaces missing values of a feature by its population mean in the test phase.

##### 3.1.4 Multiconsequence predictors

Missingness due to inapplicability could be avoided by training separate models on each molecular consequence and using only features fully applicable to each consequence. This would still leave missing values that are due to insufficient information. Missingness handling would thus still be required, even if to a lesser degree. The dimensionalities of the feature spaces of each consequence-specific classifier would be lower without a loss of information, potentially increasing performance (see also section above on reduced models). However, this requires each consequence category to have sufficiently many observations to train a classifier. In contrast, a single classifier for predicting regardless of consequence class can be trained on the whole data set, possibly allowing the predictor to generalize information between variants with different consequences.

For simplicity, we choose to focus on building a single classifier for singlenucleotide variants (SNVs) and small indels.

## 4 Materials and methods

We implement a framework that enables comparison of imputation methods by their contribution to the performance of machine-learning classifiers, specifically for prediction of variant pathogenicity. For this purpose, the framework preprocesses variant data to a usable format, performs imputation, trains a classifier, and computes relevant performance statistics.

This framework is used to perform three experiments:

1. a simulation experiment, where additional missing values are induced in the dataset several times, and the framework is used on each resulting dataset, described in subsection 4.6,
2. a cross-validation experiment, where the framework is used on datasets produced using repeated random sub-sampling, described in subsection 4.7, and
3. a main experiment, where the framework is used once with more comprehensive hyperparameter ranges for imputation methods, described in subsection 4.8.

### 4.1 Data

#### 4.1.1 ClinGen dataset

The dataset consists of ClinGen[27] expert-reviewed single-nucleotide variants from ClinVar[28, 29, 30], downloaded on 28 June 2019. We annotated the variants using the Ensembl Variant Effect Predictor (VEP)[49] version 96 with Ensembl transcripts and dbNSFP3.5 [50, 51]. We selected transcripts annotated canonical by VEP for each variant and removed the other transcripts, and where values were present for several transcripts, discarded values unrelated to the selected transcript. Any variants whose canonical VEP-annotated transcript ID did not match that from dbNSFP were discarded. In addition, we incorporated annotations used by CADD [18], matching them by transcript to the VEP-annotated Ensembl transcripts. At this stage, the dataset contained 12282 rows.

### 4.2 Preprocessing

The overall preprocessing process is depicted in Figure 1.

**Figure 1:**
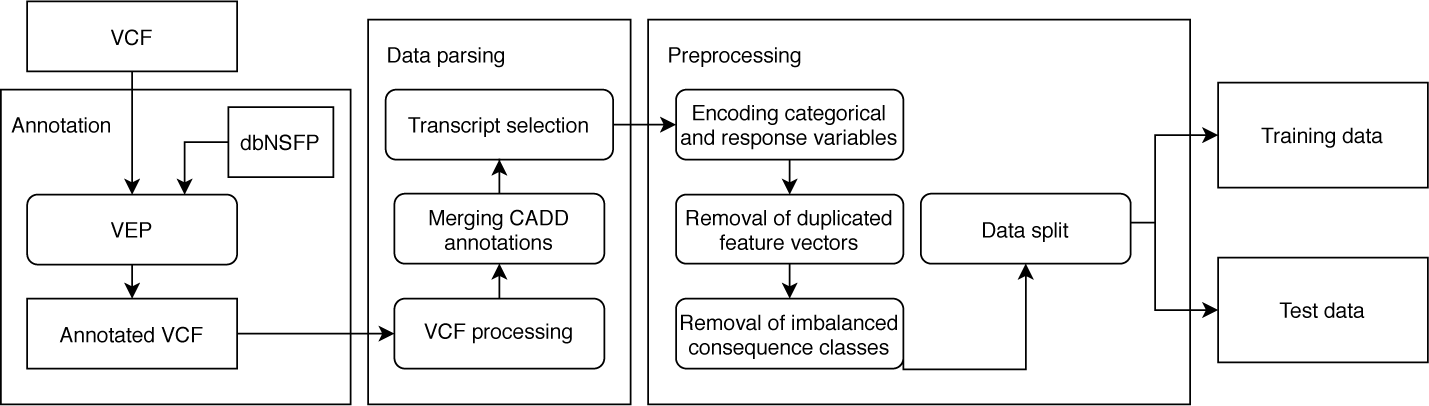
Preprocessing diagram. Data processing is divided into annotation, data parsing and preprocessing steps. Some preprocessing actions are deferred to the training phase, as their results may change due to simulated additional missing values.

The feature vectors of some sets of variants may be equal (that is, duplicated). We removed variants with duplicated feature vectors (N= 320), retaining only one variant from each equivalence class formed by duplicated feature vectors. We also chose to drop variants of uncertain significance (VUS, N = 1157) from the dataset.

The data was then randomly split into training and test subsets, with 70 % (N= 7536) of variants in the training set and 30 % (N= 3269) of variants in the test set.

We formed a binary outcome vector by defining variants classified as pathogenic or likely pathogenic to belong to the positive class (N = 5090 in the training set, N = 2218 in the test set), and variants classified as benign or likely benign to belong to the negative class (N = 2446 in the training set, N = 1051 in the test set).

Categorical features were transformed to dummy variables, with an extra category denoting a missing value. It is important to note that thus no imputation methods designed specifically for categorical features were tested, and categorical features were not treated by the used numerical imputation methods.

Use of categorical features with high class imbalance within certain levels may obfuscate the performance of the imputation methods. This is due to allowing the classifier to learn to classify all variants with that level into either the positive or the negative class, and therefore ignoring all other features upon which imputation may have been performed. VEP-predicted variant consequence is one such feature. For this reason, we removed variants with consequences for which either class had less than 5 % of overall variants of that consequence (removing 4930 variants from the training and 2139 variants from the test sets). A prediction tool developer might prefer to not make such restrictions to retain wide applicability, but it is important that they carefully analyze their results. They should especially pay attention to whether their method is significantly better than simple assignment to the majority class in such cases.

For features where the missingness implied the default value *a priori*, missing values were replaced by default values.

The final processed dataset contains 3736 variants divided into a training set with a total of 2606 variants, of which 1088 belong to the positive class and 1518 belong to the negative class, and a test set with a total of 1130 variants, of which 476 belong to the positive class and 654 belong to the negative class.

To minimize issues due to singular matrices with some imputation methods (e.g. MICE linear regression, MICE predictive mean matching), we removed features with fewer than 1 % unique values. For feature pairs with high correlation (Pearson correlation coefficient *>* 0.9), we kept only one of the features. Removal of features with few unique values or high correlation is performed as part of the training process, just before imputation, since they may be affected by introduction of additional missing values for simulations (see Simulations). It is possible that some imputation methods would be able to deal with feature sets for which the above treatment was not done, and gain an advantage due to the additional information. However, we chose to use the same restricted feature set for all imputation methods for simplicity and comparability.

A mock-up illustrating the preprocessed data format is shown in Table 2.

**Table 1:** Features imputed with default values.

| Feature name | Feature interpretation | Default value |
| --- | --- | --- |
| **motifDist** | “Reference minus alternate allele difference in nucleotide frequency within an predicted overlapping motif” [17, Supplementary information] | 0 |
| **gnomAD_exomes_AF** | gnomAD allele frequency from exomes | 0 |
| **gnomAD_genomes_AF** | gnomAD allele frequency from genomes | 0 |

**Table 2:**
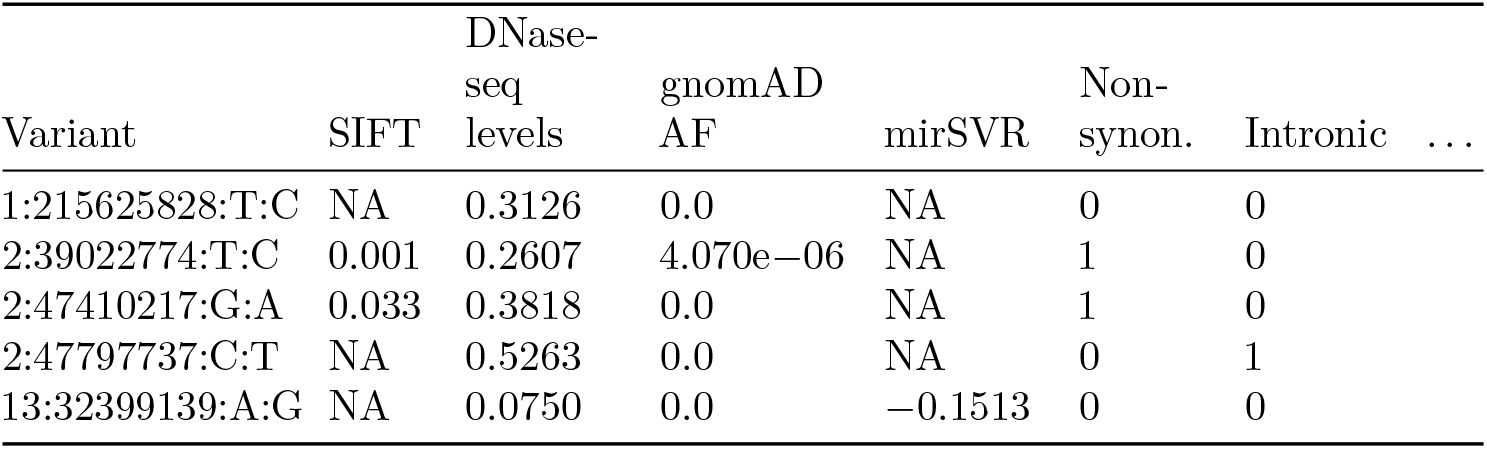
Mockup of data after preprocessing. Variant identification information is not used in imputation or training. gnomAD allele frequency has a value exactly 0 on rows 1, 3, 4, and 5 due to *a priori* imputation.

**Table 3:**
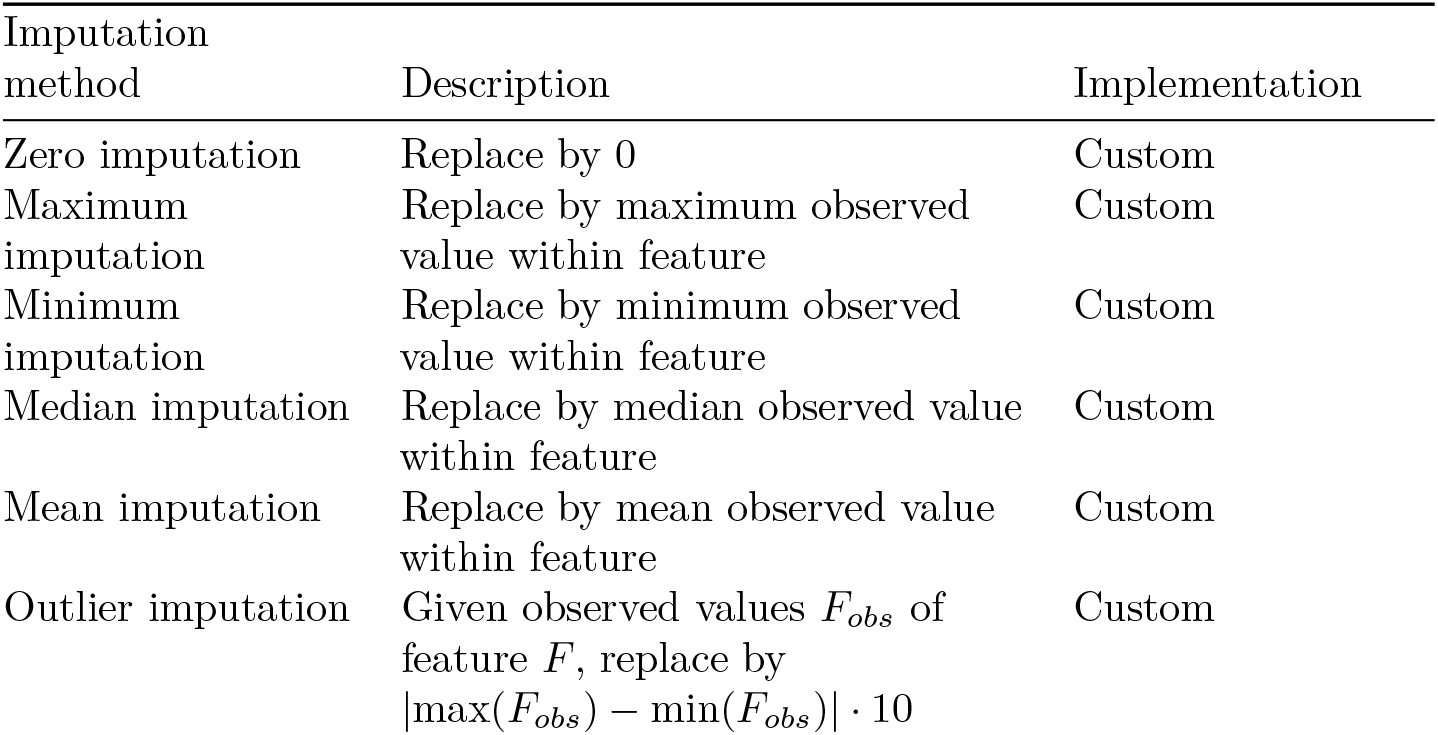
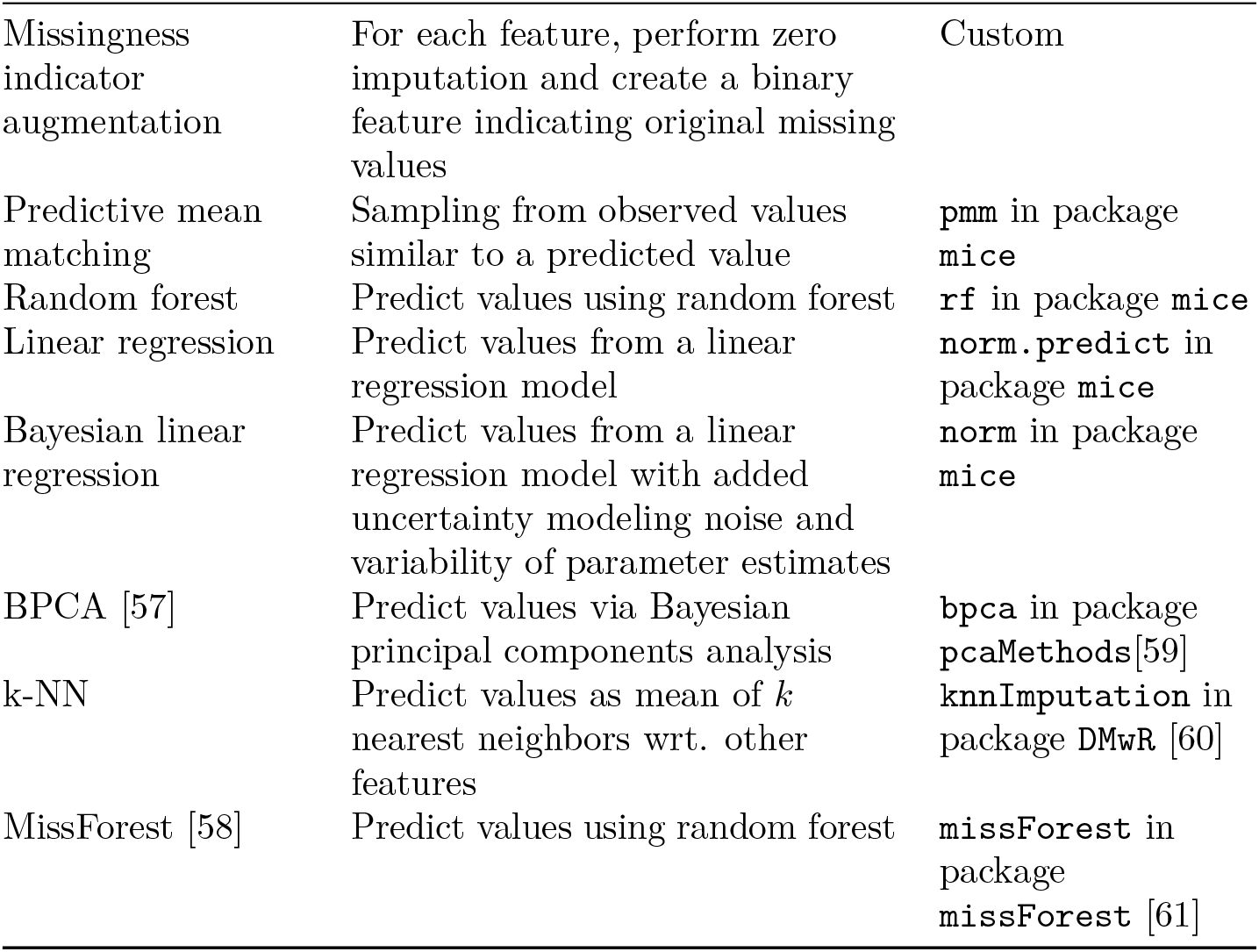
Included imputation methods

#### 4.2.1 Features

The initial feature set was defined manually to include a variety of traditional tools and annotations from both dbNSFP3.5 and the annotation set for CADD while excluding any metapredictors from the feature set. For histograms of observed values and correlations to positive outcome indicator for each feature, see supplementary information.

#### 4.2.2 Missingness

Missingness percentages of the features conditional on the VEP-predicted variant or transcript consequence are presented in Figure 2. As expected, `INTRONIC`, `UPSTREAM`, `DOWNSTREAM`, `NON-CODING_CHANGE` and `5PRIME_UTR` variants have the most missingness. They exhibit large missingness percentages across protein-related features and microRNA predictions, leaving available mainly features related to regulation. `INFRAME` variants have similar pattern as previously described variants but have observed values in features describing variant position in coding sequence and protein codon. Such features are partially observed in `SPLICE_SITE` variants, possibly due to some of them also being interpretable as coding variants. `SPLICE_SITE` variants have a feature describing the distance to a splice site completely observed. `NON-SYNONYMOUS` variants have, as expected, nearly completely observed feature vectors, except for mostly unobserved microRNA downregulation scores [52] and incompletely observed gnomAD allele frequencies, MutPred predictions and REMAP2 [53] annotations.

**Figure 2:**
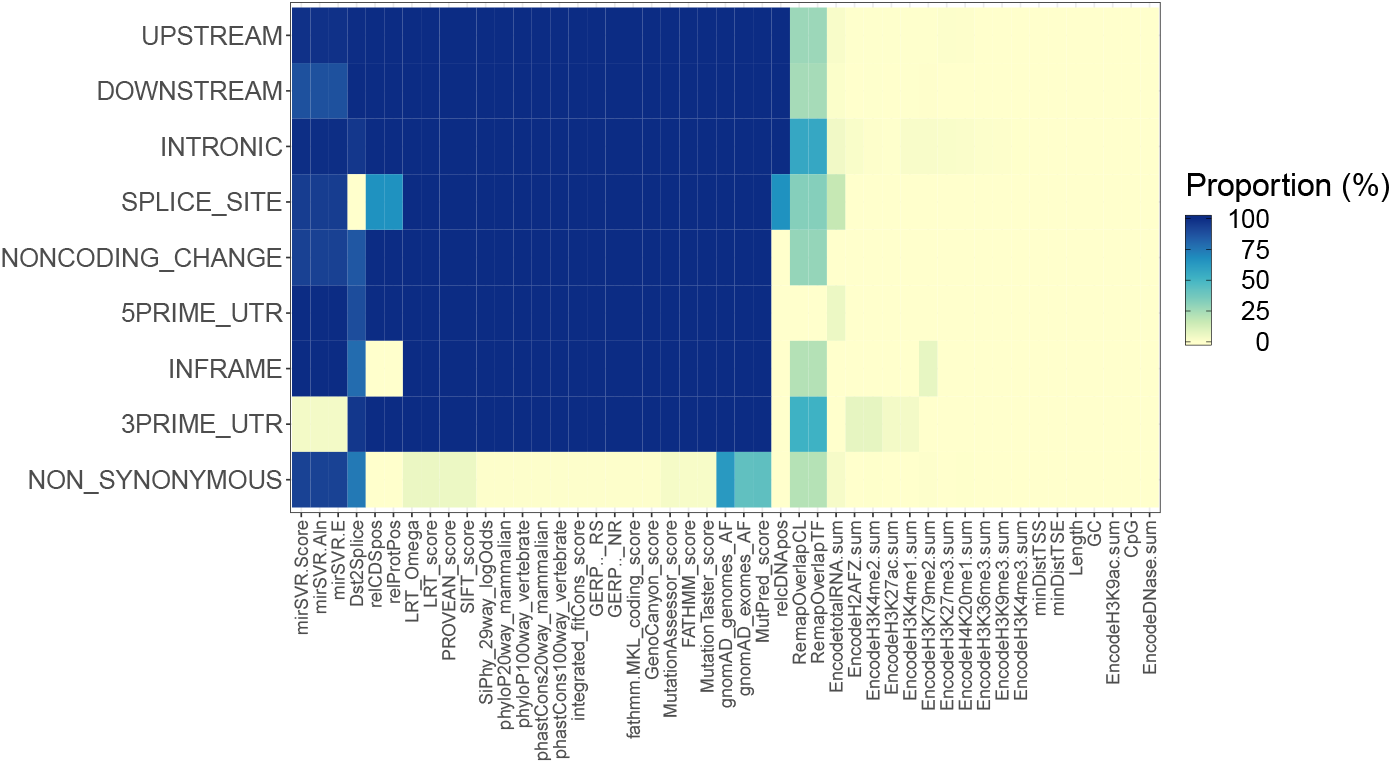
Missingness heatmap of the training set. Each cell displays the proportion of missing values of the indicated feature (horizontal axis) in the predicted molecular consequence class (vertical axis).

Since many features are only applicable to variants of a specific consequence,there will be few complete cases. Further, the only complete cases can be formed by variants that are interpretable as having several consequences, even if only one is assigned for the purposes of analysis. Such variants are, for example, splicing variants. They can often also be classified to be non-synonymous or intronic depending on their position. The complete cases are thus the very few variants that have both splicing predictions as well as the features available for non-synonymous variants.

To assess whether there is informative missingness in the data, we computed each feature’s missingness indicator’s correlation to the outcome indicator (see supplementary information). Some correlation is present for most features. Also, most correlations are negative, implying lower likelihood of pathogenicity when the feature is missing. However, correlations for `EncodetotalRNA.sum`, `gnomAD_exomes_AF` and `gnomAD_genomes_AF` are positive, implying that their missingness is linked to higher likelihood of pathogenicity.

### 4.3 Included missingness handling methods

We include, in total, 14 missingness handling methods, consisting of 6 simple imputation methods, 4 multiple imputation by chained equations (MICE) methods, 3 other popular imputation methods, and missingness indicator augmentation.

The simplest imputation methods impute every missing value within a feature with the same value, which may either be a constant or a statistic estimated from the observed values of the feature. Of these methods, we include mean, median,maximum and minimum imputation, as well as constant imputation with zero. In addition, we include outlier imputation, where we impute a feature’s missing values with a value far apart from the other values of that feature.

Multiple imputation by chained equations (MICE) [54, 55, 56] is an algorithm that iteratively imputes single features conditional on other features. In short, a MICE method

1. uses a univariate imputation method to sequentially impute each feature conditional on the observed values of other features
2. reimputes each feature conditional on the imputed data from the previous iteration

Step 2 is repeated until some maximum number of iterations or some measure of convergence is reached.

We used the R `mice` package [56] to perform the imputation. Each method was run with maximum 10 iterations.

Besides unconditional univariate methods and MICE methods, we included three popular imputation methods: k-Nearest Neighbors (k-NN), Bayesian Principal Components Analysis (BPCA) [57] and missForest [58].

For k-NN, you must have enough complete cases to start imputation, depending on *k*. As described earlier, the data had very few complete cases, and thus the largest *k* that could be used was 2.

In the case of missingness indicators, features with identical missingness patterns produce identical indicator vectors, of which only one is kept.

### 4.4 Framework

The framework performs imputation and classifier training and evaluation on a preprocessed training and test dataset pair. The preprocessed training set is used to filter out features that have insufficiently unique values or that are heavily correlated with some other feature, and this filtering is matched on the test set.

Each imputation method is used on the training set, producing at least one imputed dataset for each combination of hyperparameters (see below), and classifiers are trained on each dataset.

Multiple imputation methods can be used to produce several imputed datasets using the same hyperparameter configuration, and we make use of this in the main experiment. For probabilistic single imputation methods like MissForest, the method can be run multiple times with different seeds, producing a set of imputed datasets analogous to that generated via multiple imputation. We apply this approach to MissForest.

For each completed dataset from a probabilistic or multiple imputation method, we train a separate classifier (performing its usual hyperparameter search and model selection procedure separately on each dataset).

To maximize the performance of each imputation method for fair comparison, hyperparameter grids were defined for each method for which different hyperparameters could be passed. For the simulation and cross-validation experiments we decided to save time by using fewer hyperparameter configurations, sampling a maximum of 8 hyperparameter configurations for each imputation method used on a dataset. The hyperparameter grids are described in table 4.

**Table 4:** Statistics on hyperparameter grids for imputation methods. For k-NN, only two values of `k` succeed, see Included missingness handling methods.

| Method | Varied hyperparameters | Number of combinations |
| --- | --- | --- |
| MICE PMM | **donors, ridge, matchtype** | 180 |
| MICE regression | None | 1 |
| MICE Bayes regression | None | 1 |
| MICE Random forest | **ntree** | 36 |
| k-NN | **k** | 5 (effectively 2) |
| BPCA | **nPcs, maxSteps** | 112 |
| MissForest: | **mtry, ntree** | 12 |
| Simple methods | None | 1 each |

After imputations of the training set and classifier training, the hyperparameter configuration with highest downstream classifier performance (or highest mean performance, in case of multiple imputation methods), and its associated classifiers, are selected and stored for each method, and that hyperparameter configuration is used when imputing the test set. Finally, performance is computed using the associated classifiers and associated test set (or test sets, for probabilistic and multiple imputation) of each imputation method.

The process is depicted visually in figure 3.

**Figure 3:**
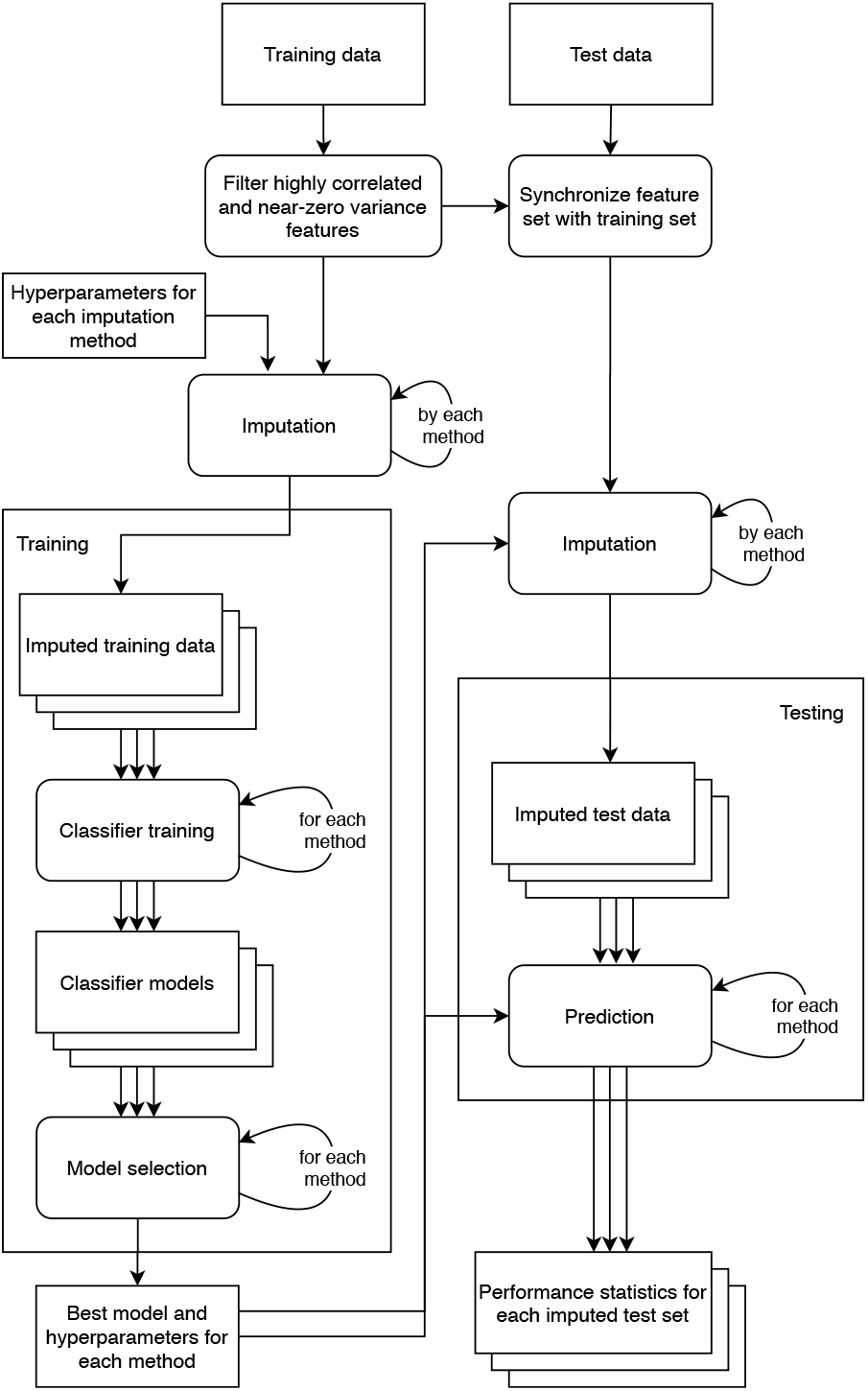
Framework flowchart. Features are filtered on the training data according to their correlations and variance, and the feature set of the test data is synchronized to match the filtered feature set. Afterwards, the training data is imputed using each imputation method, with each hyperparameter configuration. In the main experiment, multiple imputation methods are set to produce 10 imputed datasets per hyperparameter configuration. Only one dataset is set to be produced in the simulation and cross-validation experiments. Every dataset is then used to train a classifier, and for each imputation method the best-performing hyperparameter configuration is selected by the training-set performance of the corresponding classifier. For multiple imputation methods, the mean performance of the corresponding classifier set is used. The test set is then imputed with the selected hyperparameter configuration for each method, and corresponding classifier is used to predict on the imputed test set(s). Classifier performance is then evaluated on each set of predictions.

**Figure 4:**
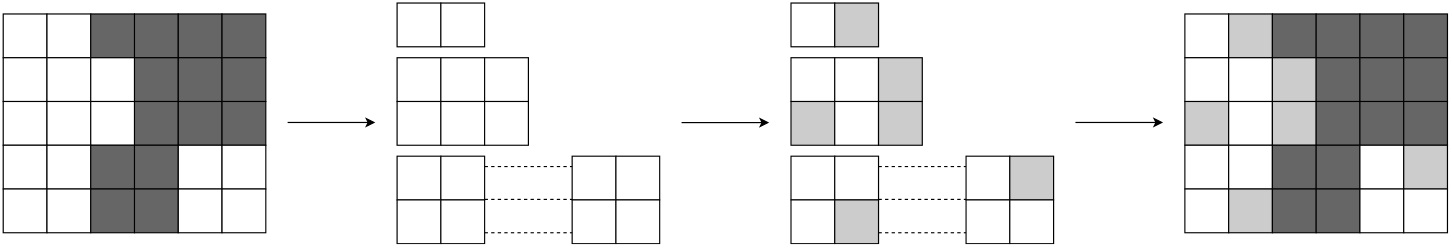
Partitioning and simulating missingness. White blocks represent available values in a data matrix, while dark gray blocks represent original missing values and light gray blocks represent additional missing values. The rows of the original dataset are partitioned to matrices with equal missingness patterns. Each feature in such matrices is now either completely observed or completely missing, and the complete features can be treated together as a fully observed matrix. Additional missingness is simulated separately using `ampute` on the fully observed matrices, which are then combined.

#### 4.4.1 Classifiers

We restricted our analysis to two common classification methods. We chose to study both a simpler, less flexible baseline method, and a more complex and highly flexible machine learning method in order to see whether classifier flexibility affects the choice of best imputation method. In this study, we refer to classifiers that are trained and predict on datasets imputed by an imputation method as *downstream classifiers* of that imputation method. For the baseline downstream classifier, we chose logistic regression (LR), a standard statistical method for two-class problems; for the more complex method, we chose random forest (RF) [38], a highly flexible machine-learning method. Both methods have been used in existing variant pathogenicity predictors (e.g. KGGSeq [47] and later versions of CADD [18] utilize logistic regression, while e.g. MutPred [62], PON-P [21], REVEL [16] and Meta-SNP [63] utilize random forest).

Logistic regression is used with the base R `glm`, and Random Forest is used via the package `randomForest` [64]. Both are trained and applied to test data via functionality from the `caret` package [65].

The random forest was trained using the out-of-bag (OOB) performance for model selection. `glm` does not offer any tuning parameters.

### 4.5 Metrics

We use *Matthews’ correlation coefficient* (MCC) as our main evaluation metric since it is less misleading than other common classification performance metrics in imbalanced datasets [66].

MCC is defined as

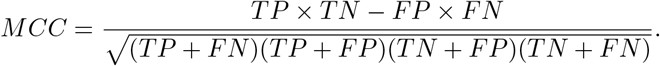

We also present results with the *area under ROC curve* (AUC-ROC, or just AUC) metric, defined as the area under the receiver operating characteristic curve.

Finally, we compute the root-mean-square error (RMSE) to compare imputed and original values when using simulated additional missing values. RMSE is defined as

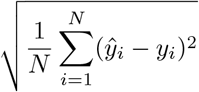

where *y*_*i*_ is the *i*th true value, 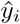 is the *i*th prediction (in this case, the imputed value) and *N* is the number of predictions (in this case, the number of imputed values).

### 4.6 Simulation

We performed simulations with additional missing values in order to

1. compare the sensitivity of imputation methods to various levels of missingness,
2. relate downstream classifier performance and an imputation method’s predictive performance, i.e. the capability of the imputation method to predict the original value underlying a missing value.

A common strategy for studying imputation methods’ performance is simulation of missing values either on fully simulated data, or on the complete subsets of real datasets. In our context of application, the number of complete cases in the dataset is very low and thus could not be used as the basis for simulating additional missing values.

Instead, we chose to take the full, incomplete dataset as the basis of simulations, and used the following strategy:

1. Create many simulated datasets based on the full dataset with additional missing values using `ampute` [67] on the dataset while varying missingness percentage
2. Impute each simulated dataset with each imputation method
3. Compute RMSE for each simulated dataset with respect to values that were observed in the original dataset but missing in the simulated dataset
4. Train a classifier on each imputed simulated training dataset, and evaluate performance on imputed original test set

We use nine percentage categories of additional missingness, from 10 % to 90 %, each producing 100 datasets with the respective amount of additional missingness. We therefore have 900 simulated datasets on which the framework is executed. To reduce the computational requirements, we downsample the hyperparameter grids of each imputation method (in comparison to the main experiment). In addition, since MissForest is significantly more computationally expensive than other imputation methods, it is not included in the simulation experiment.

#### 4.6.1 Amputing additional missing values

The input matrix of `ampute` is required to be complete, so we partitioned the data according to missingness patterns. This forms a set of matrices for which every feature is either fully observed or fully missing. We then used each of the fully observed submatrices as inputs to `ampute`, and combined the resulting output back to form a matrix of the original size.

#### 4.6.2 Simulation evaluation

Many imputation studies are specifically built to assess an imputation method’s capability to predict the original values from the observed values of a dataset with missing data. Thus, they use the RMSE between the original dataset and an imputed dataset as a metric of performance of the imputation method. However, an imputation method with the lowest RMSE is not in general the best one in the statistical inference context [32, chapter 2.6]. In short, the best imputation method wrt. RMSE is linear regression estimated via least squares, and the deterministic nature of a regression prediction necessarily ignores uncertainty due to the missing data. The same argument cannot be applied to the predictive context, but the same end result may apply when missingness is informative.

Consider a situation where missingness is highly informative. Then a perfect imputation method (with respect to RMSE; i.e. one where RMSE would equal 0) would impute the dataset with the original values. However, the informativeness in the missing values would be lost. The loss of information could, in principle, be avoided by adding missingness indicators before imputing, but this comes with the increase in dimensionality (doubling the number of features in the worst case) and thus cannot be seen as a universal solution.

Since RMSE cannot be used as a universal metric of imputation method performance, we perform a lean version of the framework on each simulated dataset. That is, we impute each simulated dataset using a sparser hyperparameter grid (downsampled to eight hyperparameter configurations separately for each simulated dataset) and producing only one dataset each when using probabilistic methods. A classifier of each type is trained on the imputed simulated datasets, the best performing imputation hyperparameter configuration is chosen by the highest performing classifier trained on a dataset imputed via that configuration. Performance is estimated on the test set imputed with the winning configuration. We compute, in addition, RMSE of each imputation method on each dataset, and can thus investigate whether low RMSE on the training set predicts high downstream classifier performance on the test set.

As mentioned in the Preprocessing section, due to the removal of features with fewer than 1 % unique values and features that highly correlate with another feature, the addition of missing values may lead to differing feature sets between different simulated datasets. Especially large numbers of additional (MCAR) missing values may lead to fewer unique values, and both increase or decrease correlation between features by chance.

### 4.7 Cross-validation experiment

In the cross-validation experiment, the training data is used to produce 100 training/test dataset pairs via repeated random sub-sampling with a 70 % split. The framework is run separately on each pair, after which the results can be used to estimate imputation methods’ relative robustness to variation from sampling. For speed, hyperparameters for imputation methods are downsampled to eight hyperparameter configurations each, and multiple imputation methods are set to produce only a single dataset. However, due to the fewer required executions of the framework (when compared to the simulations), it was feasible to also include MissForest in the set of methods.

### 4.8 Main experiment

In the main experiment, the framework is run once with the full hyperparameter grid for each imputation method on the full training and test datasets, and MissForest is included in the set of methods. We also utilize the multiple datasets generated by multiple imputation methods to estimate the performance variability that arises from randomness in the imputation of both training and test sets.

### 4.9 Challenges

#### 4.9.1 Dataset composition

Even after *a-priori* imputation, there is a very high proportion of missing values in the dataset (∼ 31 %). The missingness exhibits a general pattern, and is not e.g. monotone (see [31, pp. 6–7]). In addition, missingness appears in most features, and the number of complete cases ends up minuscule (2 rows in training data), and removal of any single feature would not significantly increase the number of available complete cases (see figure 2). The missingness can also not in general be assumed to be MCAR, though due to the arguments by Sarle [33] and Ding & Simonoff [34] mentioned earlier, we expect this has little effect on classification performance.

#### 4.9.2 Studying imputation methods developed for statistical inference in a predictive context

As mentioned in subsection 3.1.1, the differences in intended usage between statistical inference and prediction make it difficult to use existing implementations of imputation methods–many of which were designed for the former purpose–in prediction. The first and main issue is that out-of-the-box implementations often do not provide an easy way to reuse learned parameters from an earlier run. This makes it difficult to use parameters from the training set on the test set.

Some ways to deal with this are

- *Reimplementation*. One can reimplement the method in a way that allows reuse of parameters, fully solving the problem. However, this is work-intensive, and requires deep understanding of each imputation method.
- *Ignore*. One can ignore the issue, and allow the imputation method to re-estimate its parameters when imputing the test set. However, this may lead to diminished classifier performance on the test set, as the distributions of imputed values may differ between training and test sets and thus may confound the classifier.
- *Concatenation of incomplete datasets*. When imputing the test set, concatenate the training set and test set, impute the combined dataset and then remove rows belonging to the training set. This reduces the difference between the distributions compared to *Ignore*, but may be computationally expensive.
- *Concatenation to imputed training data*. One can make a variation on *Concatenation of incomplete datasets* by imputing the training set before concatenation. This is faster than *Concatenation of incomplete datasets* if the imputation method does not run in linear time with respect to the size of the input dataset. However, it may introduce additional issues, since it will use the imputed values in the training set to estimate imputations for the test set.
- *Concatenation of single observation*: A second variation of *Concatenation of incomplete datasets* is to concatenate only one observation from the test set at a time, and repeating this until the full test set is imputed. This variation makes it possible to predict on a single observation at a time. This is computationally expensive.

*Reimplementation* and *Concatenation of single observation* are the only options that allow imputation of test observations independently from each other. The others perform imputation with parameters estimated from other observations that are being predicted on at the same time.

| Imputation method / family | Implementation | Out-of-the-box parameter reuse |
| --- | --- | --- |
| MICE | `mice` | No |
| BPCA | `pcaMethods` | No |
| k-NN | `DMwR` | Yes |
| Simple methods | `custom` | Yes |
| MissForest | `missForest` | No |

The package `mlr`[68] offers wrapper functionality that allows use of any prediction method offered by the package also for univariate imputation, along with functionality for correct reimputing data with previously learned parameters, but using this option is difficult in a dataset with very few complete cases. We did not explore this possibility in this work. Investigation of the imputation performance of methods originally intended for prediction might merit further study.

We choose to implement option *Ignore* due to its simplicity for methods where out-of-the-box parameter reuse is not available. There is a possibility that this will give an advantage to simple methods and k-NN even if MICE, BPCA and MissForest would otherwise outperform them. However, this comparison still represents the situation as it presents itself to the practitioner that may not have the time or expertise to make use of the other options.

### 4.10 Implementation

The framework and experiments were implemented with R [26] and organized into 11 executable scripts:

| Number | Description | File name |
| --- | --- | --- |
| 1. | Parsing of VCF file | 01_parse_vcf.R |
| 2. | Preprocessing | 02_preprocess_data.R |
| 3. | Computation of descriptive statistics | 03_descriptive_stats.R |
| 4. | Execution of training set imputation and classifier training for main experiment | 04_run_impute_and_train.R |
| 5. | Execution of test set imputation and prediction for main experiment | 05_run_prediction.R |
| 6. | Generation of datasets with additional missing values | 06_generate_simulated_data.R |
| 7. | Execution of imputation and classifier training on datasets with additional missing values | 07_run_simulations.R |
| 8. | Process and plot test set performance statistics for main experiment | 08_analyze_results.R |
| 9. | Process test set performance statistics for classifiers trained on datasets with additional missing values | 09_analyze_simulation_results.R |
| 10. | Plot test set performance statistics for classifiers trained on datasets with additional missing values | 10_simulations_plots.R |
| 11. | Perform repeated random sub-sampling cross-validation and process and plot its results | 11_crossvalidation.R |

The scripts are intended to be executed in order, but users may choose only run a subset if they are only interested in a subset of the results. To run the main experiment, one must run 1., 2., 4., 5., and 8.; to run the simulations, one must run 1., 2., 6., 7., 9. and 10.; to run the cross-validation experiment, one must run 1., 2. and 11.

### 4.11 Availability of source code and requirements

Project name: AMISS

Project home page: https://github.com/blueprint-genetics/amiss

Operating system(s): Linux

Programming language: R [26]

Other requirements: The software was run with R 3.6.0 with packages vcfR [69], futile.logger [70], tidyr [71], here [72], magrittr [73], ggcorrplot [74], mice[56], foreach [75], doParallel [76], ggplot2 [77], iterators [78], missForest [61, 58],DMwR [60], doRNG [79], rngtools [80], lattice [81], itertools [82], randomForest[64], ModelMetrics [83], stringr [84], gridExtra [85], digest [86], purrr [87], caret[65, 88], testthat [89] and e1071 [90], and pcaMethods [59] via BioConductor [91]BiocManager [92].

License: MIT

Any restrictions to use by non-academics: CADD annotations require commercial users to contact authors for licensing. dbNSFP [51] annotations may require licenses for commercial use and must be reviewed individually.

### 4.12 Availability of supporting data and materials

The result files from experiments described in this article are available in the Zenodo repository, DOI 10.5281/zenodo.6656616 [93].

## 5 Results

This section is arranged as follows. First, we present results from the added-missingness simulations. We evaluate both the relation of downstream classifier performance and RMSE, and how increasing missingness percentage affects downstream classifier performance. Next, we present results from the repeated random sub-sampling cross-validation experiment, shedding light on robustness of missingness handling methods to dataset composition. We then present results pertaining the main experiment, comparing the downstream classifier performances of missingness handling methods. Finally, we compare the missingness handling methods with respect to their running times.

### 5.1 Simulation experiments

#### 5.1.1 RMSE

We found that RMSE and classification performance as measured by MCC did not correlate in datasets with simulated additional missing values (for plots of RMSE against MCC for each imputation method, see supplementary information). This is especially clear in the case of outlier imputation, where RMSE is–as expected–much higher than with any other method, while MCC was comparable to other methods.

#### 5.1.2 Missingness percentage

The effect of additional MCAR missingness on MCC performance of downstream classifiers is displayed in figures 5 and 6. Fitted LOESS (locally estimated scatterplot smoothing) curves are shown.

**Figure 5:**
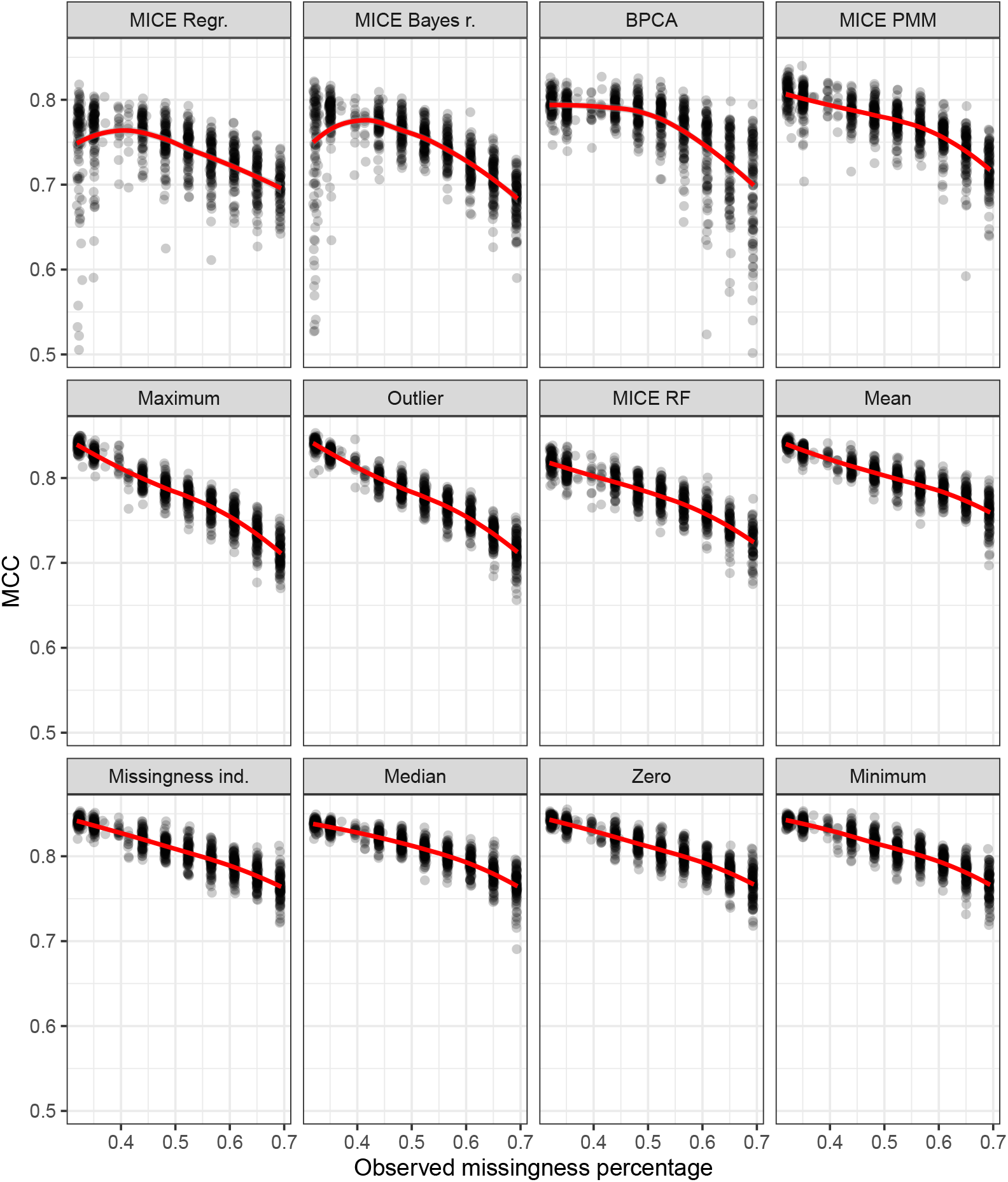
Random forest MCC against actual missingness percentage, with fitted LOESS curves (red).

**Figure 6:**
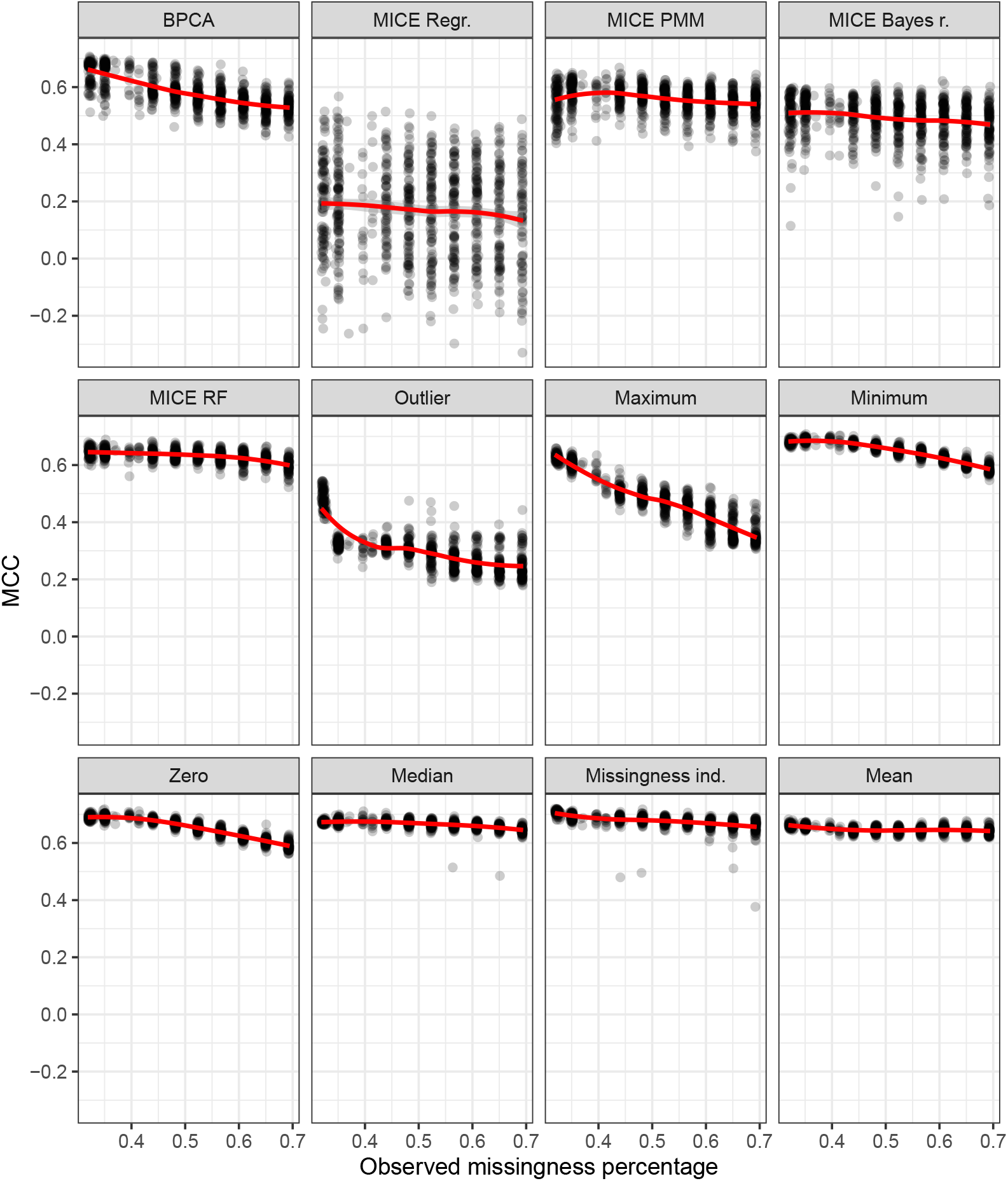
Logistic regression MCC against actual missingness percentage, with fitted LOESS curves (red).

When the downstream classifier is a random forest, missingness indicator augmentation as well as mean, minimum, zero and median imputations show similar curves, with their average performances dropping from slightly below MCC = 0.85 at 30 % missingness to slightly above MCC = 0.75 at 70 % missingness. MICE random forest and MICE PMM have overall slightly lower mean performance than the previously mentioned methods, but otherwise show similar shapes. Outlier and maximum imputations suffer more drastically, with mean performances dropping to just above MCC = 0.70 at 70 % missingness. MICE regression and MICE Bayes regression demonstrate a curious effect where average downstream classifier performance increases at first as missingness increases, before starting their descent. BPCA shows hints of a similar but more muted trend. This may be related to a phenomenon noted by Poulos & Valle [94] in the context of prediction on categorical variables, where introduction of additional missing values prior to imputation may improve classifier performance.

When the downstream classifier is logistic regression, outlier imputation shows a dramatic drop immediately between 30 % and 40 % missingness and stabilises slightly above MCC = 0.20. Maximum imputation shows a clear linear downward trend, while minimum imputation, zero imputation and BPCA show a much smaller one. Missingness augmentation and all MICE methods show very light reductions in average performance as missing value percentage grows. Median and especially mean imputation show practically no performance reduction due to increasing MCAR missingness. MICE Bayes regression, BPCA and PMM and especially MICE regression all show much larger variability in their performance than other methods.

### 5.2 Cross-validation experiment

The variability of downstream classifier performance evaluated via repeated random sub-sampling cross-validation is displayed in figure 7.

**Figure 7:**
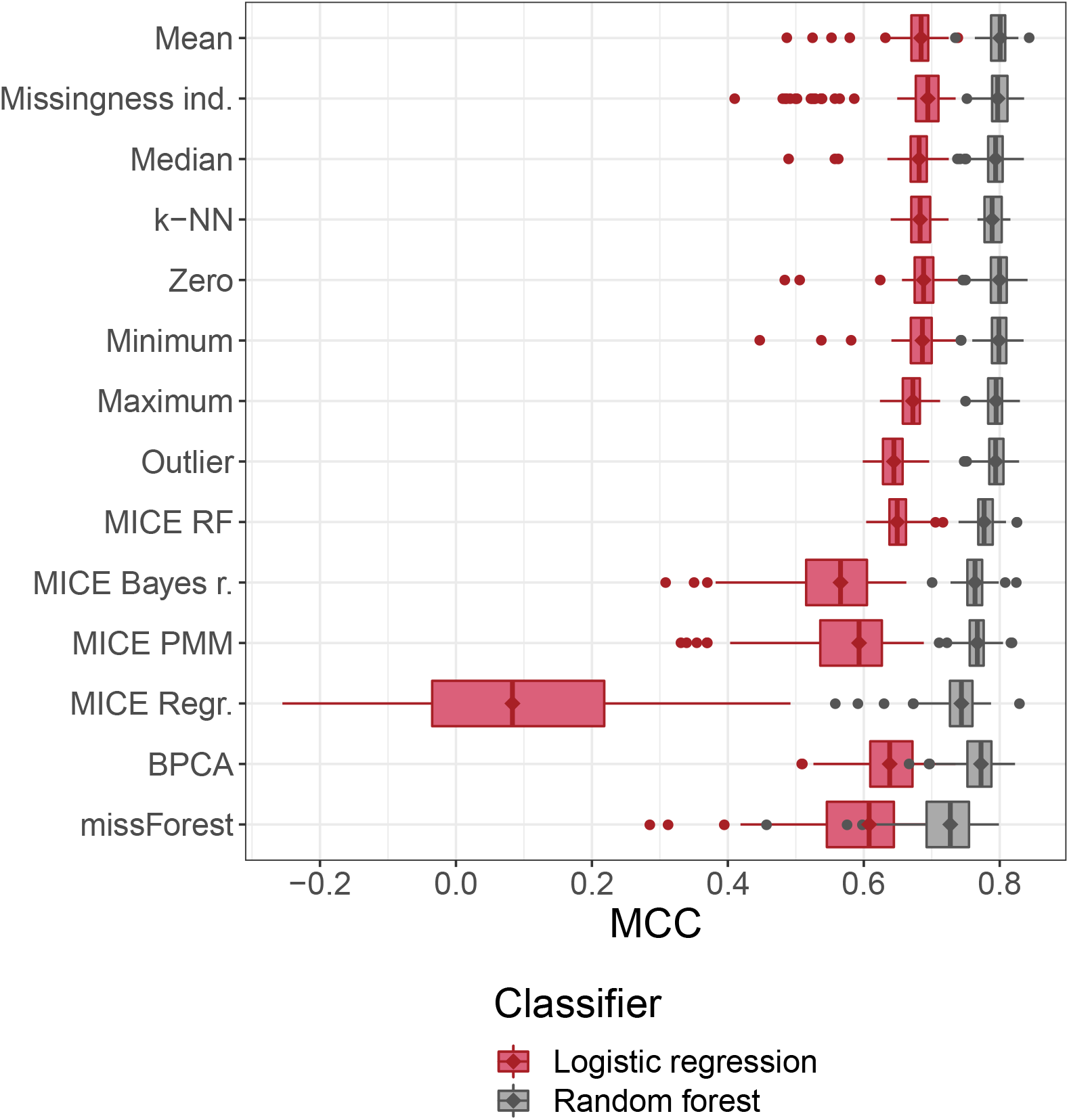
Boxplots describing down-stream classifier performance estimated via repeated random sub-sampling cross-validation. Diamonds were added to emphasize median values. The training set was further randomly split 100 times into 70%*/*30% subsets, and the framework was executed for each of the resulting pairs.

The random forest classifier outperforms logistic regression regardless of imputation method, and the lowest performing imputation method wrt. random forest classifiers (MissForest) has higher mean performance than the highest performing imputation method wrt. logistic regression (missingness indicators).

For both logistic regression and random forest, single imputation methods andk-NN appear to have equivalent performance, though in conjunction with logistic regression outlier imputation seems to perform slightly worse. With regard to random forest classifiers, MICE random forest, BPCA, MICE Bayes regression, and MICE PMM perform slightly or somewhat worse than simple imputation methods, but display greater differences in conjunction with logistic regression, where MICE random forest and BPCA are clearly preferable to MICE Bayes regression and MICE PMM. MICE ordinary regression and MissForest perform worse on average with both classifier types, but when combined with logistic regression, MICE ordinary regression brings mean classification performance down close to that of a coin flip.

### 5.3 Main experiment

In order to assess the differences in downstream classifier performance due to selection of imputation method, we show mean performance metrics for classifiers trained on datasets produced by each missingness handling method, sorted by MCC, for random forest classifiers (table 7) and for logistic regression classifiers (table 8). Single imputation methods (mean, missingness indicator, median, k-NN, zero, minimum, maximum, outlier) produced only a single dataset, and as such also produced only a single set of performance statistics. MICE ordinary regression imputation was unable to produce imputations in the main experiment, and is thus excluded.

**Table 7:** Mean test set performance metrics for a random forest classifier trained and tested on data sets treated with each missingness handling method

| Method | MCC | AUC-ROC | Sensitivity | Specificity | $F_1$ | Precision |
| --- | --- | --- | --- | --- | --- | --- |
| Maximum | 0.848 | 0.975 | 0.912 | 0.937 | 0.913 | 0.914 |
| Missingness indicators | 0.846 | 0.975 | 0.908 | 0.937 | 0.910 | 0.913 |
| k-NN imputation | 0.844 | 0.973 | 0.918 | 0.930 | 0.911 | 0.905 |
| Mean | 0.844 | 0.974 | 0.914 | 0.934 | 0.912 | 0.910 |
| Zero | 0.840 | 0.976 | 0.905 | 0.933 | 0.906 | 0.907 |
| Median | 0.839 | 0.975 | 0.916 | 0.925 | 0.907 | 0.899 |
| Minimum | 0.838 | 0.975 | 0.903 | 0.934 | 0.906 | 0.909 |
| Outlier | 0.837 | 0.974 | 0.908 | 0.930 | 0.906 | 0.904 |
| MICE RF | 0.817 | 0.976 | 0.889 | 0.927 | 0.894 | 0.898 |
| MICE Bayes regression | 0.812 | 0.975 | 0.876 | 0.932 | 0.890 | 0.904 |
| MICE PMM | 0.812 | 0.975 | 0.869 | 0.936 | 0.888 | 0.909 |
| BPCA | 0.781 | 0.974 | 0.779 | 0.976 | 0.860 | 0.959 |
| MissForest | 0.756 | 0.952 | 0.818 | 0.927 | 0.853 | 0.892 |

**Table 8:** Mean test-set performance metrics for a logistic regression classifier trained and tested on data sets treated with each missingness handling method

| Method | MCC | AUC-ROC | Sensitivity | Specificity | $F_1$ | Precision |
| --- | --- | --- | --- | --- | --- | --- |
| Missingness indicators | 0.700 | 0.925 | 0.840 | 0.862 | 0.828 | 0.816 |
| k-NN | 0.682 | 0.924 | 0.817 | 0.865 | 0.816 | 0.816 |
| BPCA | 0.680 | 0.923 | 0.828 | 0.855 | 0.817 | 0.806 |
| Mean | 0.674 | 0.920 | 0.815 | 0.859 | 0.812 | 0.808 |
| Zero | 0.673 | 0.923 | 0.809 | 0.864 | 0.811 | 0.812 |
| Minimum | 0.672 | 0.922 | 0.813 | 0.859 | 0.810 | 0.808 |
| Maximum | 0.670 | 0.914 | 0.828 | 0.846 | 0.812 | 0.796 |
| Outlier | 0.670 | 0.906 | 0.863 | 0.813 | 0.815 | 0.771 |
| Median | 0.669 | 0.921 | 0.815 | 0.855 | 0.809 | 0.803 |
| MissForest | 0.645 | 0.929 | 0.695 | 0.921 | 0.769 | 0.868 |
| MICE RF | 0.642 | 0.909 | 0.761 | 0.874 | 0.787 | 0.815 |
| MICE PMM | 0.601 | 0.898 | 0.709 | 0.874 | 0.751 | 0.810 |
| MICE Bayes regression | 0.591 | 0.879 | 0.785 | 0.805 | 0.763 | 0.750 |

In the case of random forest classification, maximum imputation has the best average performance with respect to MCC (= 0.848). It is very closely followed by missingness indicator augmentation, k-NN imputation and mean imputation, and then zero, median, minimum and outlier imputation methods. MICE random forest imputation is the highest performing MICE method (MCC = 0.817), with slightly better MCC than Bayes regression and PMM. The final group, with distinctly worse downstream classifier MCC, consists of BPCA (MCC = 0.781) and MissForest (MCC = 0.756).

Mean classification performance is lower across the board for logistic regression. Missingness indicator augmentation is the winner in this scenario with MCC = 0.700. k-NN imputation (MCC = 0.682) and BPCA (MCC = 0.680)receive second and third place, after which mean imputation, zero imputation, minimum imputation, maximum imputation, outlier and median imputations have essentially equal performance (MCC ranging from 0.674 to 0.669). MissForest and MICE methods (MCC between 0.645 and 0.591) perform noticeably worse.

The MCC and AUC-ROC performances of downstream classifiers are illustrated in figures 8 and 9, respectively. For multiple imputation and MissForest the variability of performance due to randomness in the imputation is also visible. Variability in downstream classifier performance is seen to be smaller for random forest compared to logistic regression in all methods except for MissForest, and variability in general larger in methods with lower mean performance.

**Figure 8:**
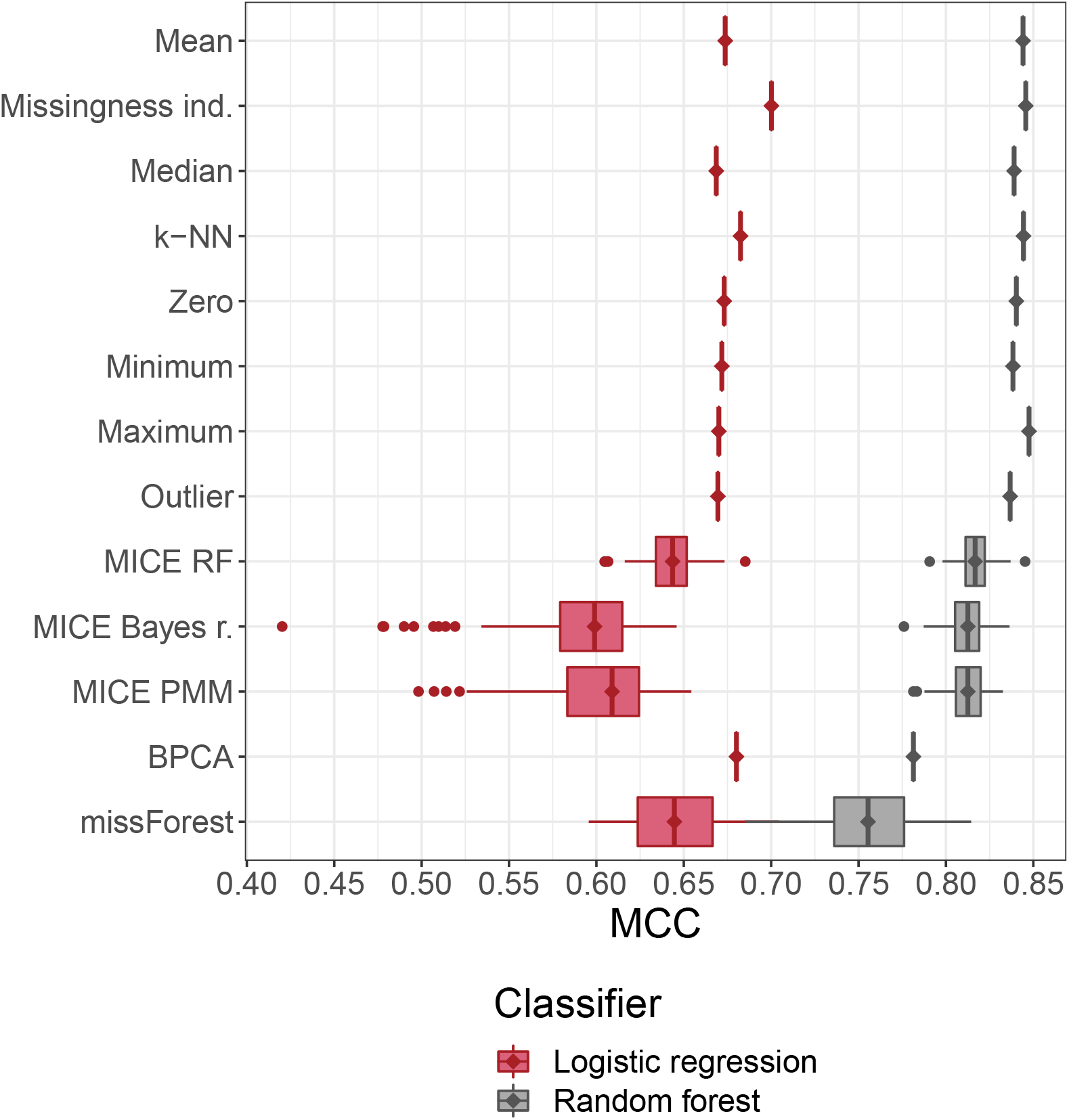
Boxplots describing down-stream classifier performance with respect to MCC. Diamonds were added to emphasize median values. Variance represents variation in probabilistic or multiple imputation, with 10 classifiers, each trained on a training set imputed with the same method, used to predict on 10 test sets imputed using the same method.

**Figure 9:**
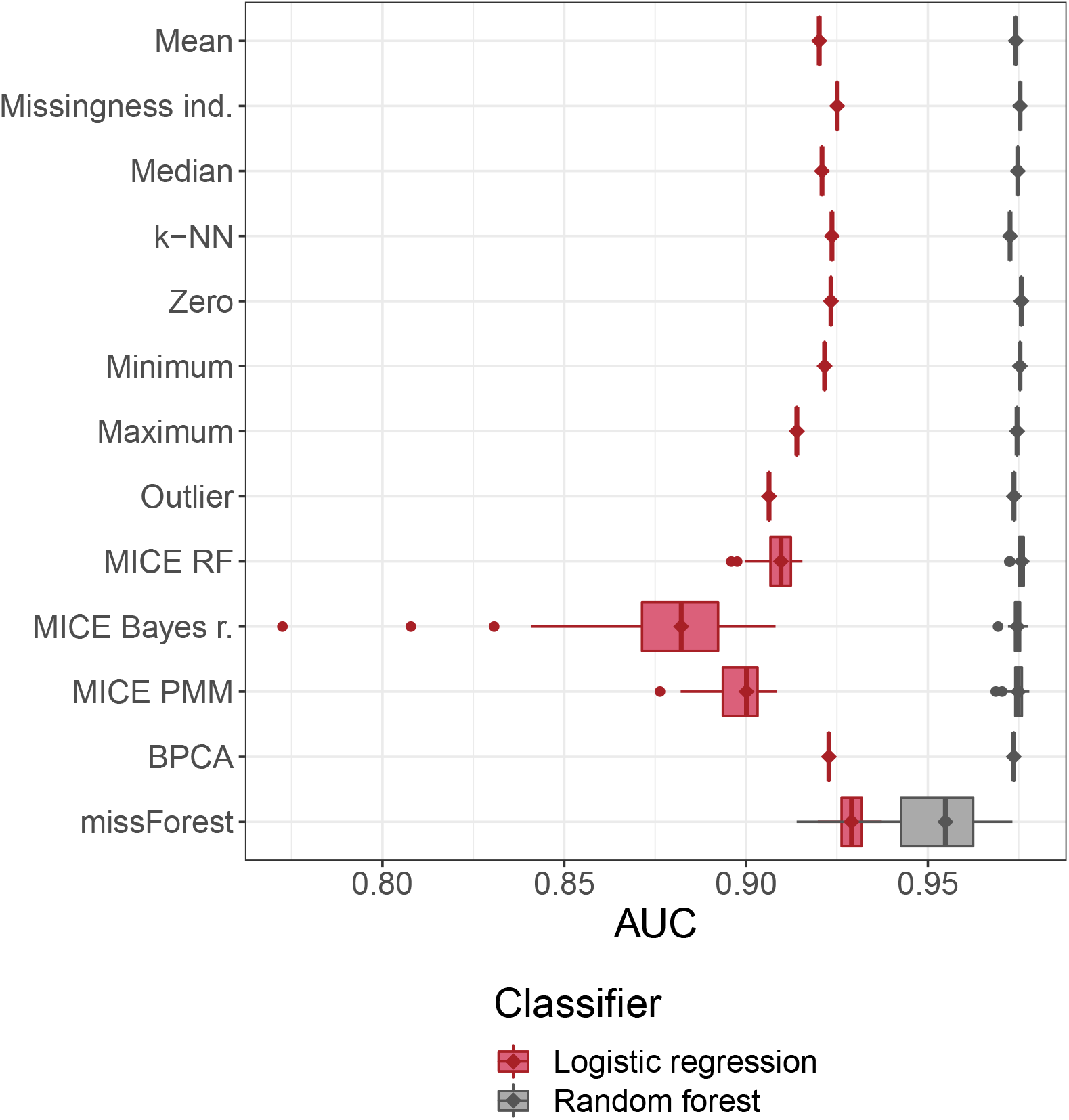
Boxplots describing down-stream classifier performance with respect to AUC-ROC. Diamonds were added to emphasize median values. Variance represents variation in probabilistic or multiple imputation, with 10 classifiers, each trained on a training set imputed with the same method, used to predict on 10 test sets imputed using the same method.

Using AUC-ROC to measure performance makes it more difficult to compare imputation methods due to the very small absolute differences. For a randomforest classifier (table 7), the mean AUC-ROC of all methods, with the exception of MissForest, is within 0.002 of each other. For logistic regression (table 8), the range is wider, and is also visually distinguishable in figure 9. Interestingly, here MissForest has the upper hand. However, looking back at table 8 we notice that MissForest shows the highest specificity but also the lowest sensitivity of all methods. The imbalance is reflected in the relatively poor MCC and *F*_1_ scores, but ignored by AUC-ROC.

The results are not very different from the cross-validation experiment results. When compared using MCC, simple imputation methods are largely interchangeable, though missingness indicators dominate when using a logistic regression classifier, and maximum imputation is dominant when using a random forest classifier. However, when considering the variance shown in the cross-validation experiment, the differences between simple imputation methods are likely to be due to chance. It is important to note that here the variance for multiple imputation methods is due to the multiple imputation (or multiple runs of a probabilistic imputation method) of training and test sets, and thus is not directly comparable to that of the cross-validation experiments. MICE ordinary regression is not included here due to it failing in the main experiment.

#### 5.3.1 Results grouped by consequence

To assess whether the strongly distinct missingness patterns exhibited within different variant consequence classes, we also computed performance statistics separately within certain consequence classes, specifically `DOWNSTREAM`, `UPSTREAM`, `INTRONIC`, and `Other`, an aggregate of all remaining consequence classes. `Other` is formed by the consequence classes for which either 1) there are overall few variants, and the class indicator gets filtered out due to non-zero variance, or 2) there is high correlation with another feature. Results are shown in figure 10.

**Figure 10:**
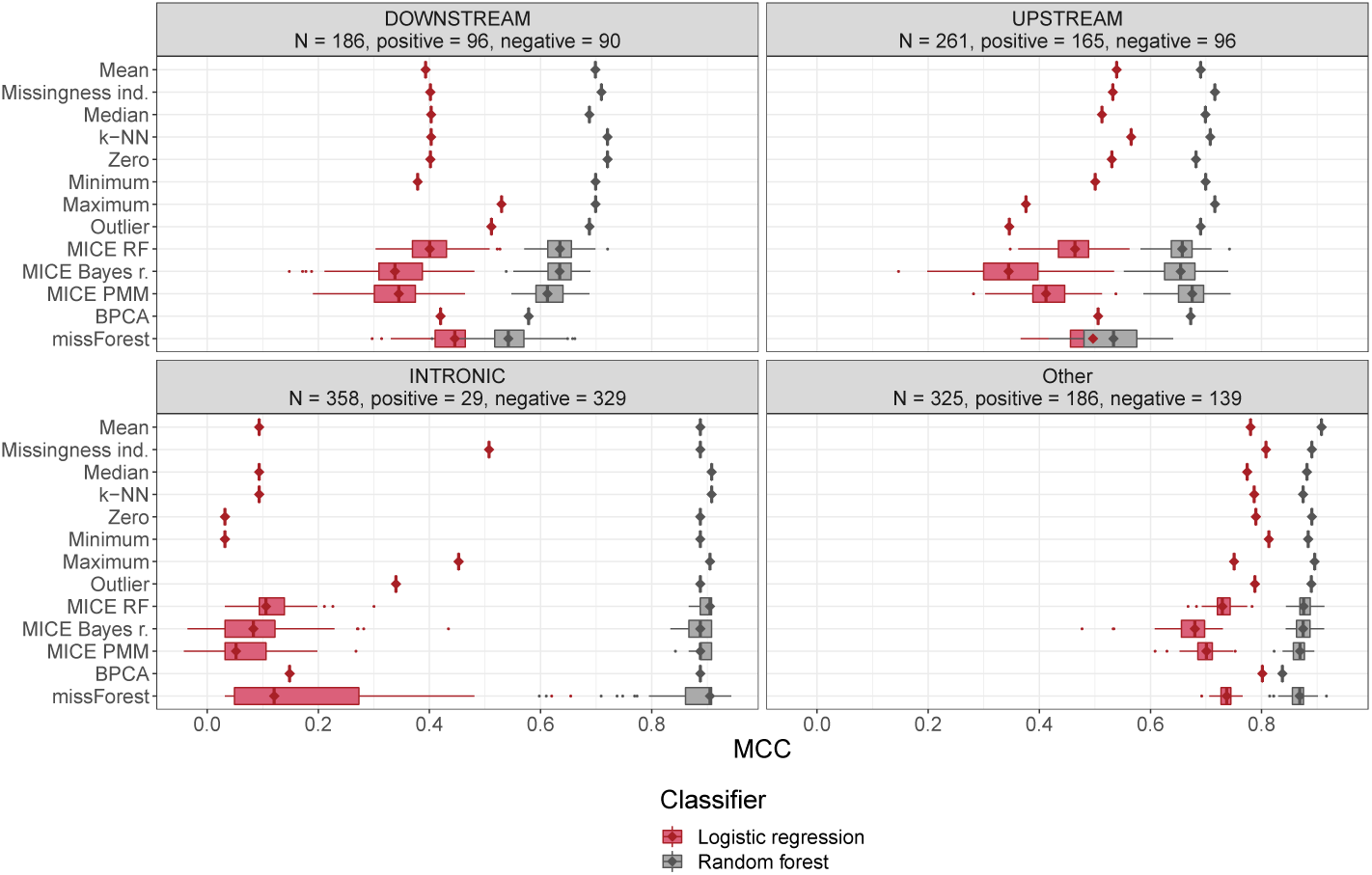
Boxplots describing down-stream classifier performance with respect to MCC, conditional on variant consequence. Diamonds were added to emphasize median values. Variance represents variation in probabilistic or multiple imputation, with 10 classifiers, each trained on a training set imputed with the same method, used to predict on 10 test sets imputed using the same method.

Some consequence classes were indeed very sparse, as seen in table 9, and the pre-imputation filtering process eliminated the dummy variables encoding classmembership of `INFRAME`, `NONCODING_CHANGE`, `5PRIME_UTR`, `3PRIME_UTR`, and`SPLICE_SITE`. In addition, membership to class `NON_SYNONYMOUS` was strongly (negatively) correlated to the missingness of LRT predictions, and was also eliminated by the pre-imputation feature filtering. Thus these classes were grouped into `Other`.

**Table 9:** Numbers of occurrence of variant consequence classes.

| Consequence | Occurrences |
| --- | --- |
| INFRAME | 14 |
| NONCODING_CHANGE | 14 |
| 5PRIME_UTR | 18 |
| 3PRIME_UTR | 27 |
| SPLICE_SITE | 53 |
| DOWNSTREAM | 464 |
| NON_SYNONYMOUS | 578 |
| UPSTREAM | 632 |
| INTRONIC | 806 |

Performances show large differences in different consequence classes. `Other` is mostly non-synonymous variants, as described above, and shows good performance for both classifier types, though random forest is always superior. Mean imputation performs best with the random forest classifier, while missingness indicator augmentation and minimum imputation maximize logistic regression performance. The results are essentially equal to the general results depicted in figure 8 where consequence classes are not distinguished.

The consequence class `INTRONIC` shows a very different and interesting situation. First it is important to note that even after the preprocessing steps which removed variants in consequence classes with high class imbalance, only *≈*8.1 % of `INTRONIC` variants are positive. However, the robustness of MCC to class imbalance allows us to compare the performances. In this consequence class, logistic regression is virtually useless, though missingness indicator augmentation and maximum imputation seem to allow some discriminatory power. It seemsthat in `INTRONIC`, logistic regression learns to classify almost everything as negative (see supplementary information for plots of sensitivity and specificity conditional on consequence class). With most imputation methods, MCC is barely above zero. In contrast, the random forest classifier performs well with any imputation method, though the high variation in MissForest imputations seems to allow also for instances of poor performance.

Compared to `Other`, prediction on variants of `DOWNSTREAM` consequence shows lower performance in both classifiers. For random forest, the relative orders of imputation methods are basically the same as in the main experiment and the cross-validation experiment. Unlike in all other consequence classes, logistic regression seems to perform better with MICE methods and MissForest than most simple imputation methods and k-NN, though maximum and outlier imputation do seem to outperform them even here.

In `UPSTREAM` variants, performance is again lower than in `Other`, but not as low as in `DOWNSTREAM`. Random forest classification shows the common pattern, where simple imputation methods and k-NN seem slightly preferable to MICE methods, and MissForest performs noticeably worse. Logistic regression classification looks, in this case, to be disadvantaged by the same methods that dominated `DOWNSTREAM` (that is, maximum and outlier imputation).

As mentioned earlier, minor differences between simple imputation methods have a good chance of being attributable to randomness.

### 5.4 Running time

Running times were recorded in the main experiment for the best hyperparameter configurations of all imputation methods and are presented in table 10. For methods that were used to produce multiple datasets (MICE methods and MissForest) the overall time was recorded and then divided by the number of produced datasets (10). The machine used to run the software was a CentOS Linux server with two Intel Xeon CPU E5-2650 v4 @ 2.20GHz processors and 512 GiB of memory. The analysis was run with parallelization using 24 processes. Each imputation was run inside a single process, and thus running times should not be affected based on whether an imputation method itself offers parallelization features.

**Table 10:** Running time for imputing the training set with the best hyperparameter configurations in the main experiment.

| Method | LR (elapsed (s)) | RF (elapsed (s)) |
| --- | --- | --- |
| Zero | 0.006 | 0.006 |
| Maximum | 0.007 | 0.007 |
| Minimum | 0.007 | 0.007 |
| Mean | 0.009 | 0.009 |
| Median | 0.011 | 0.011 |
| Outlier | 0.014 | 0.014 |
| Missingness indicator augmentation | 0.478 | 0.478 |
| MICE Bayes regression | 5.379 | 5.379 |
| MICE regression | 5.839 | 5.839 |
| k-NN | 12.792 | 12.948 |
| BPCA | 14.884 | 19.511 |
| MICE PMM | 15.427 | 14.921 |
| MICE RF | 107.036 | 104.278 |
| MissForest | 362.352 | 362.352 |

MICE regression and MICE Bayes regression show similar running times and finish between 5 and 6 seconds, while k-NN, MICE PMM and BPCA take somewhat longer. MICE random forest and especially MissForest take much longer. Simple imputation methods are much faster than all other methods.

## 6 Discussion

Random forest classifiers defeated logistic regression overall, with the worst-performing combination of missingness handling method and random forest still reaching higher performance than best-performing combination of missingness handling method and logistic regression. This was expected, due to both the inherent capability of tree predictors such as those in random forest to ignore irrelevant features, and the more flexible decision boundary that random forests are able to form. Choice of missingness handling method thus cannot compensate for an unsuitable classification method.

The high comparative performance of k-NN with both classifiers is surprising. The depletion of complete cases greatly limits the possible values for *k*. Even more importantly, the few complete cases are the only possible neighbors to any point; when *k* equals the number of complete cases, every missing value in a specific feature will be imputed with the average value of that feature in the set of complete cases. k-NN imputation with *k* equaling the number of complete cases can thus be interpreted to be a form of unconditional imputation, since in such a case every missing value will be imputed with a single value, irrespective of the values of other features.

The computational cost of different methods varies dramatically. Many of the highest-performing methods take a minuscule portion of time spent during the overall training and prediction process, while more costly methods dominate the time requirements. The difference between zero imputation (0.006 second runtime on our dataset) to MissForest (362.352 seconds) may be inconsequential for a single sample, but is compounded with large cohorts or high-throughput diagnostic work. For example, for 100 000 WGS samples the difference in total computational time with the simplest and most complex method will be over ayear, potentially translating into hundreds of thousands of euros of additional cloud computing and storage costs.

Simple imputation methods overall could be considered the winning group of the comparison presented in this paper, but mean imputation might further be pointed out as the favorite in that group. In addition to the consistently high MCC in all experiments, it also displays high resilience to performance degradation due to increasing MCAR missingness, and low variance in relation to sampling. It is also one of the fastest methods to compute, trivial to implement and is always applicable.

Considering the complexity of the prediction task, the features, the large degree of missingness, and the sophistication of available missingness handling strategies, it is somewhat surprising that the best performance is gained using simple unconditional imputation strategies.

### 6.1 Possible explanations to simple imputation superiority

#### 6.1.1 Informativeness

A putative explanation is the presence of some informativeness in the missingness mechanism. If the presence of a missing value in certain variables correlates with the pathogenicity of the variant, simple imputation methods would be given an advantage: when every imputed value (within a given feature) is replaced by a single specific value, the classifier may learn to correlate that single value with the outcome. This is less likely to happen when using more sophisticated imputation methods, which make it harder for the classifier to learn which values were likely imputed. However, logistic regression is less flexible and thus less capable of representing such potentially discontinuous dependencies, and yet missingness indicator augmentation did not perform dramatically better than most simple imputation methods. This would imply that informativeness does not hold a large role, even though we did find it to be present in the data. It may also be that the dimensionality increase due to the additional indicator features disadvantages the classifier enough to neutralize the benefit from informativeness.

#### 6.1.2 Inability to transfer training parameters to test set

As described in Methods and materials, many available implementations of imputation methods do not allow reuse of parameters between imputation of training and test sets. Thus the estimated distributions on which imputed values are based will differ at least slightly, and thus disadvantage any methods for which parameter reuse was not possible.

### 6.2 Additional challenges

Grimm et al. [95] described several biasing factors in variant effect predictor training and performance evaluation using data from commonly used variant databases, e.g. the tendency for variants within the same gene being classified as all pathogenic or all neutral, or simply due to difficulty of finding datasets completely disjoint with the training set. Mahmood et al. [96] further analysed existing variant effect predictors using datasets generated from functional assays, and found drastically lower performance compared to earlier reported estimates.

Our approach is not immune to these biases, and we expect that any reported performance metrics will be overoptimistic. However, we expect that the main result of the study, the relative performance rankings of missingness handling methods, is not affected by the biases. The classifiers built described in this work are not intended to outperform earlier approaches or be directly used for variant effect prediction.

### 6.3 Conclusions

It appears that it is unnecessary to use sophisticated missingness handling methods to treat missing values when building variant pathogenicity metapredictors. Instead, simple unconditional imputation methods and even zero imputation give higher performance and save significant computational time, leading to considerable cost savings if adopted. This highlights the conceptual separation between missingness handling methods for prediction and imputation for statistical inference, the latter of which requires carefully constructed techniques to reach correct conclusions.

### 6.4 Further work

There are several ways to improve and expand on this work. The dataset could be extended to include variants from wider sources, and the effect of circularity could be estimated using additional datasets. It would likely also be possible to make changes to or reimplement methods whose implementation does not currently support reuse of parameters. Reduced models and its hybrid variants would make an interesting point of comparison if implemented. Another possible extension of the work would be to broaden the focus from missingness handling to various other design choices that may affect predictor performance, such as using random search in place of grid search or downsampling data to improve class balance.

## 7 Potential implications

We compared a variety of commonly used missingness handling methods in order to assess their suitability for using machine learning to build variant pathogenicity metapredictors. The analysis will help pathogenicity predictor researchers choose missingness handling methods to maximize the performance of their tools.

## 9 Author contributions

JP conceived and supervised the study. MS performed the study and prepared the manuscript. SM performed testing and code review. SM, IS, KM, VF, LL and JP reviewed and approved the manuscript.

## 10 Acknowledgements

Not applicable

## 11 Consent for publication

Not applicable

