## Supplementary information for "Comparison of missing data handling methods for variant pathogenicity predictors"

Figures 1 and 2 show downstream classifier performance in the simulation experiment, plotted against RMSE across columns for a subset of the imputation methods, plotted with equal scales for each method. Outlier imputation is excluded here since its RMSE values are much higher than other methods. As can be seen in the figure, RMSE and MCC are not correlated, especially well illustrated by maximum, minimum and zero imputation, where high MCC is reached even as RMSE is consistently higher than for other methods.

Figures 3 and 4 are as above, except with only the MCC scale fixed. Outlier imputation and BPCA have drastically higher RMSE, but outlier imputation still shows good MCC values with a random forest classifier.

Figures 5 and 6 show the sensitivity and specificity, respectively, for test set variants of different consequence classes in the main experiment. Logistic regression seems to have little discriminatory power in `INTRONIC` variants when combined with most missingness handling methods, as sensitivity is near zero, implying all variants in this consequence class are being classified as benign. Missingness indicators, outlier imputation and maximum imputation seem to allow the method to better detect pathogenic variants, though still at a lower rate than random forest.

Figure 7 shows correlations in the training data from feature values to the positive outcome indicator.

Figure 8 shows correlations in the training data from the missingness indicators of each feature to the positive outcome indicator. Consequence.x, CpG, EncodeD-

NAse.sum, EncodeH3K9ac.sum, GC, Length, minDistTSE and minDistTSS have no missing values and thus have an empty correlation bar. For most features, the correlation is negative, implying that missingness in those features decreases the likelihood of pathogenicity. Correlations from missingness indicators of EncodetotalRNA.sum, gnomAD_exomes_AF and gnomAD_genomes_AF are positive, implying that missingness in those features increases the likelihood of pathogenicity.

Figures 9 through 16 show the histograms of feature values for each feature on the training set.

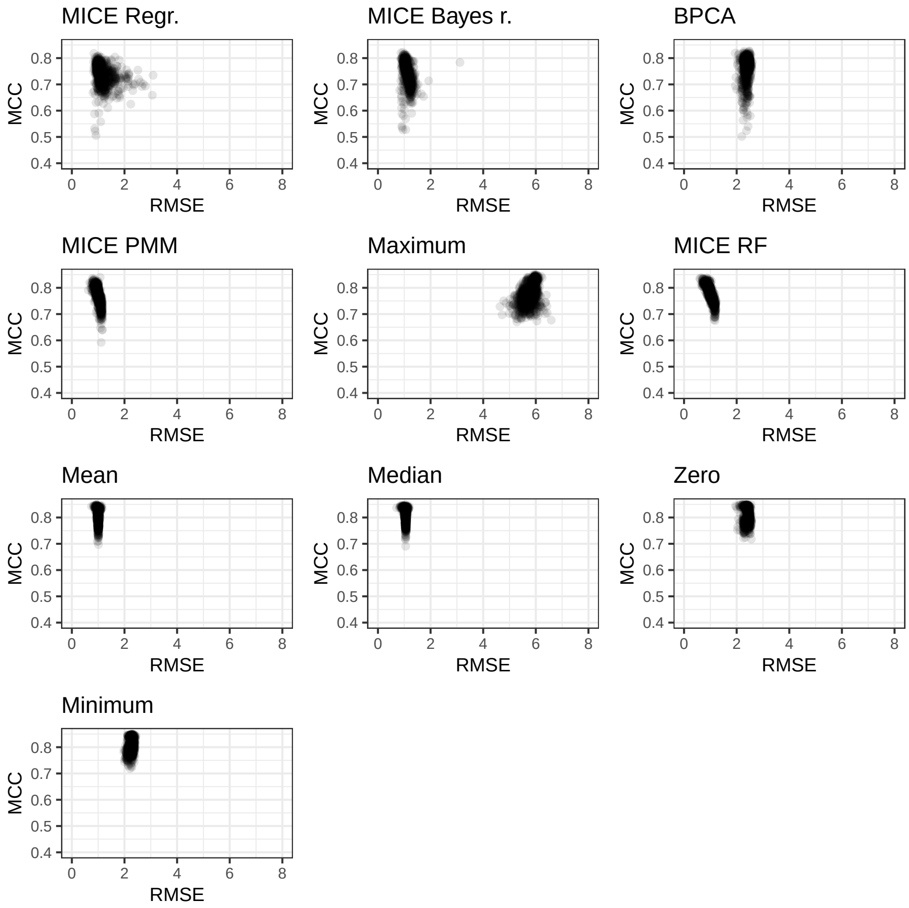


Figure 1: Random forest MCC against RMSE averaged across columns, equal scales

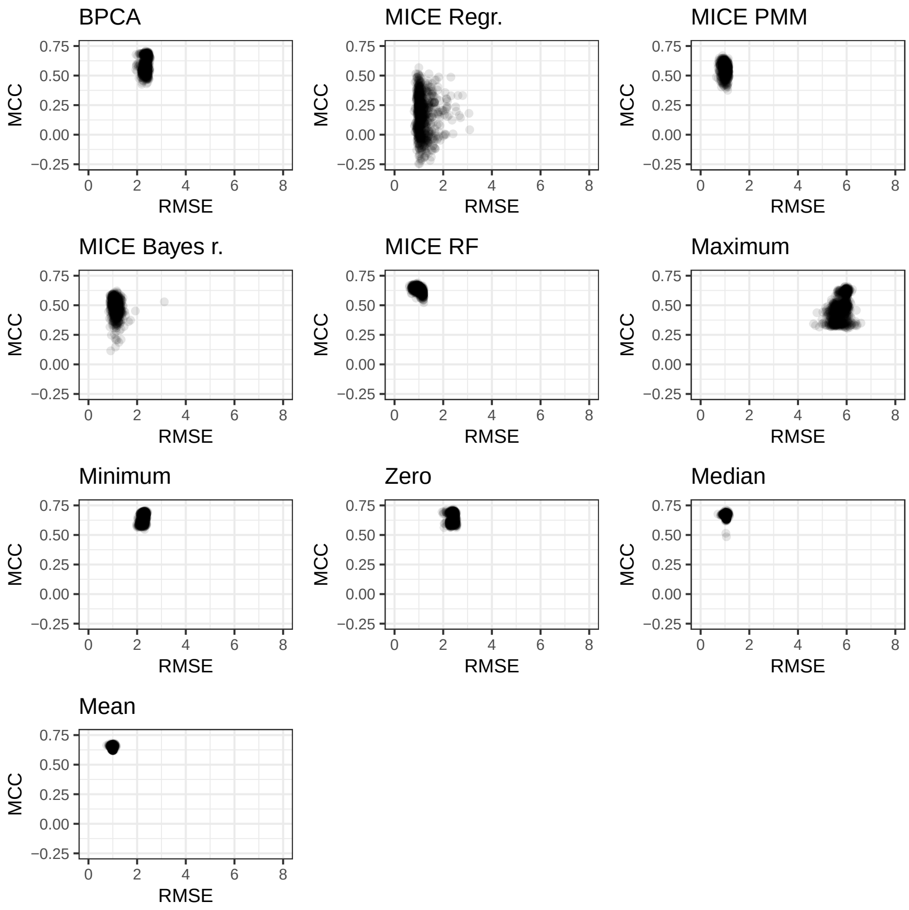


Figure 2: Logistic regression MCC against RMSE averaged across columns, equal scales

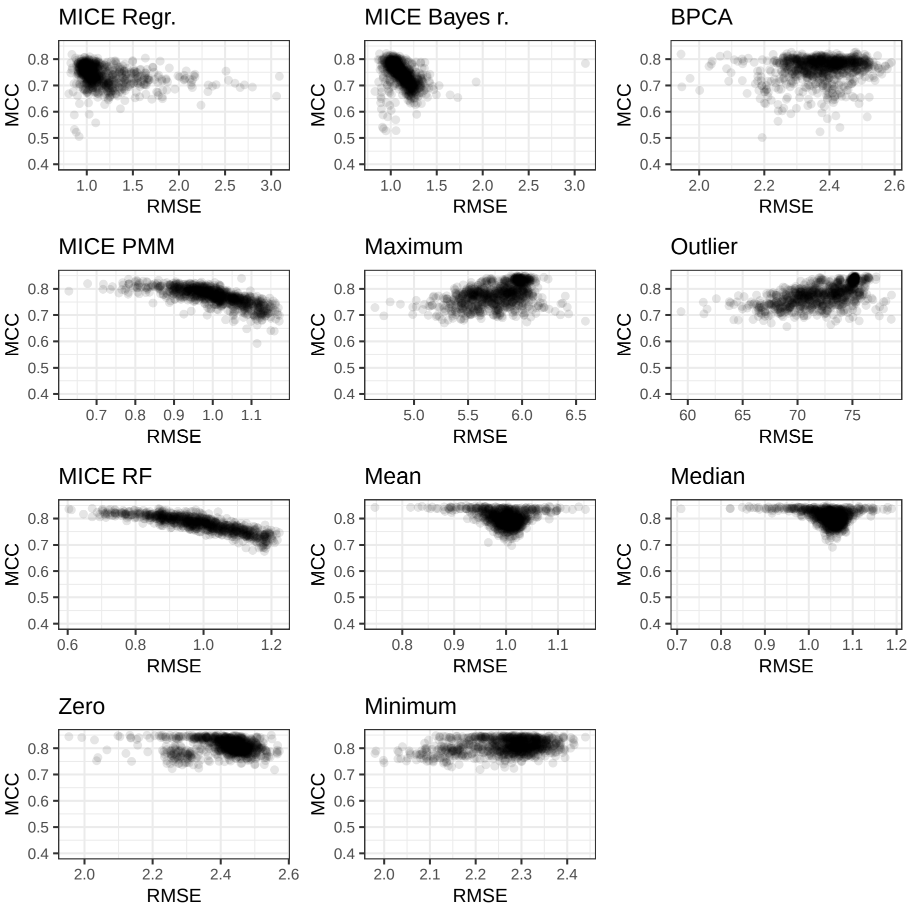


Figure 3: Random forest MCC against RMSE averaged across columns, free `x` scale

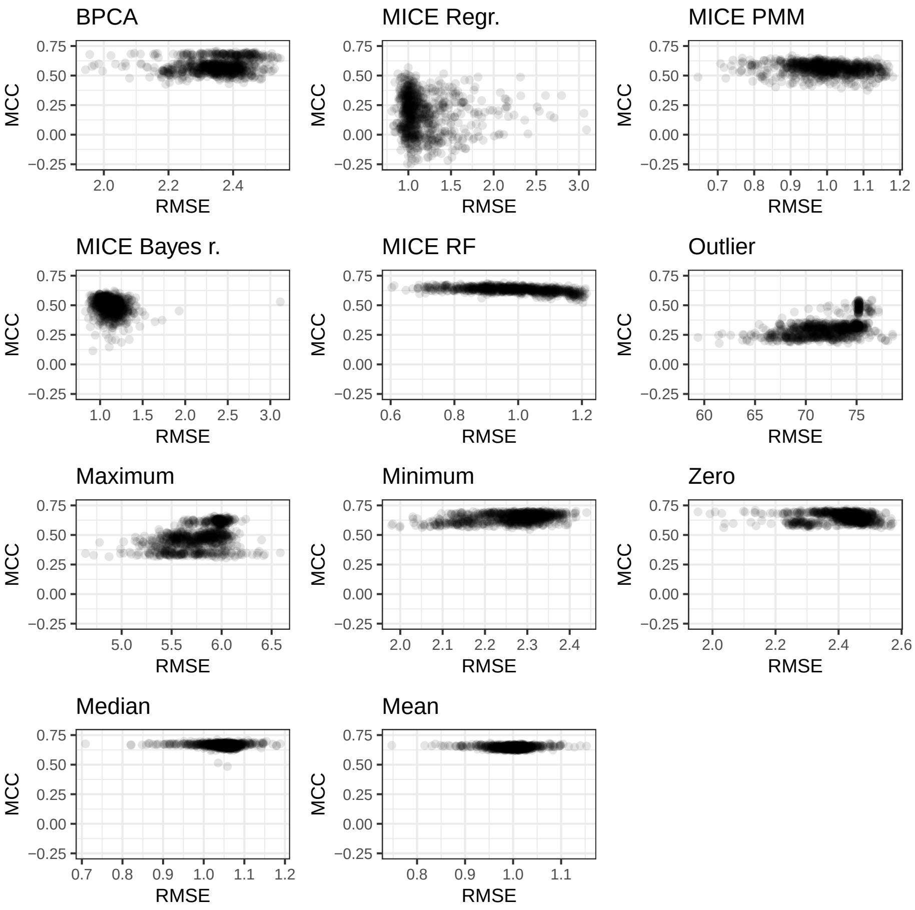


Figure 4: Logistic regression MCC against RMSE averaged across columns, free `x` scale

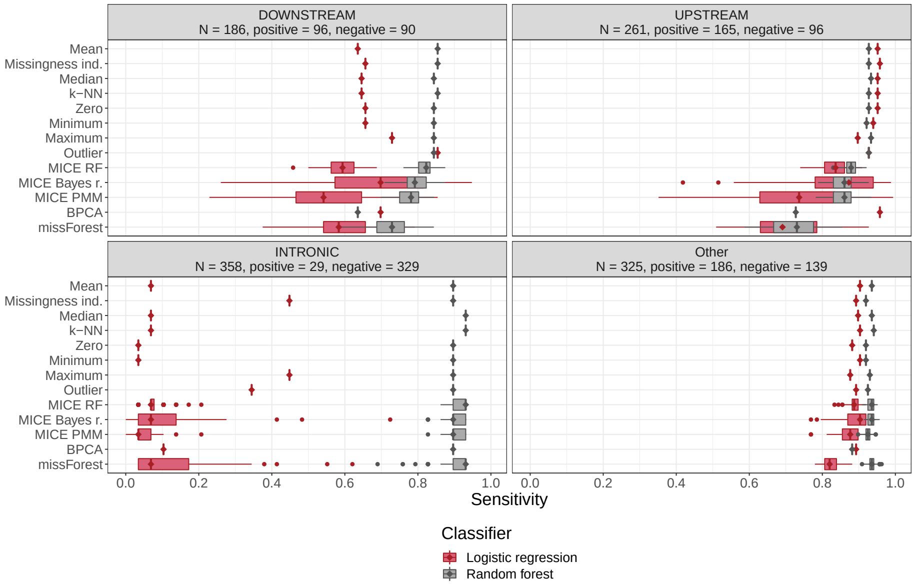


Figure 5: Sensitivity boxplots for both RF and LR classifiers, conditional on variant consequence.

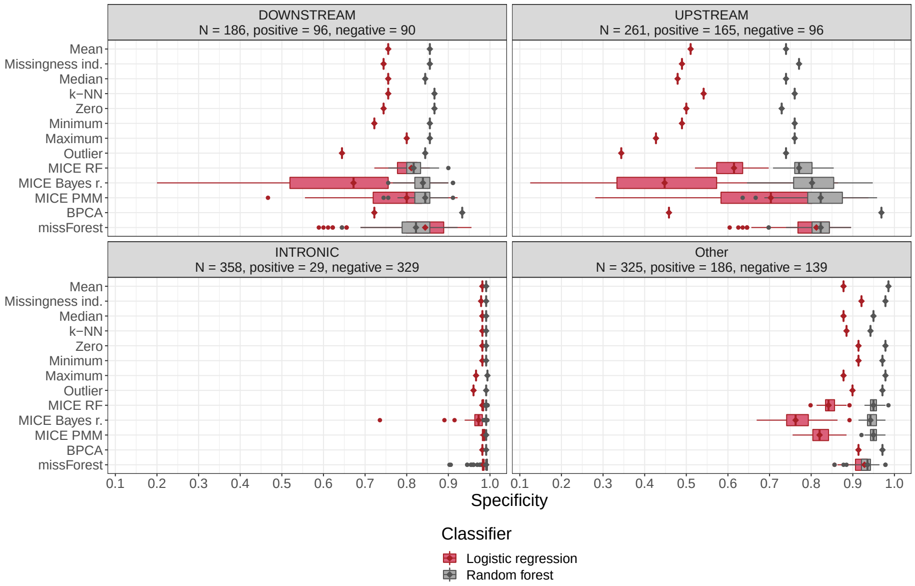


Figure 6: Specificity boxplots for both RF and LR classifiers, conditional on variant consequence.

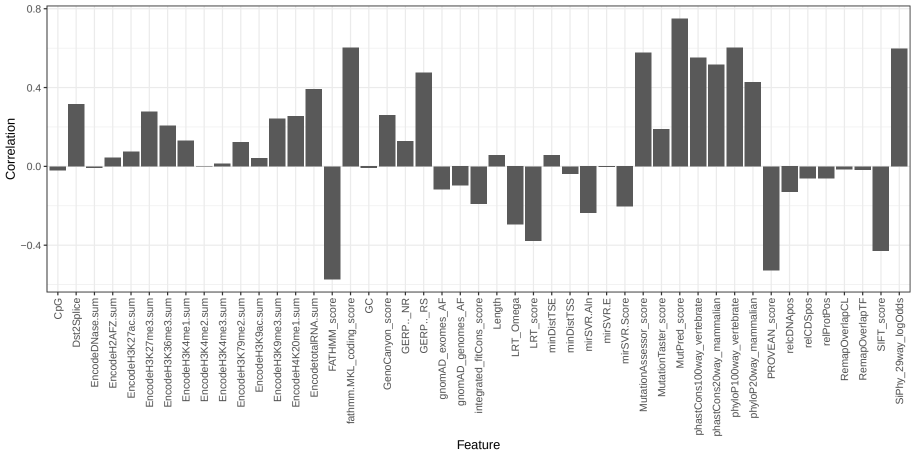


Figure 7: Correlations of features to positive outcome indicator.

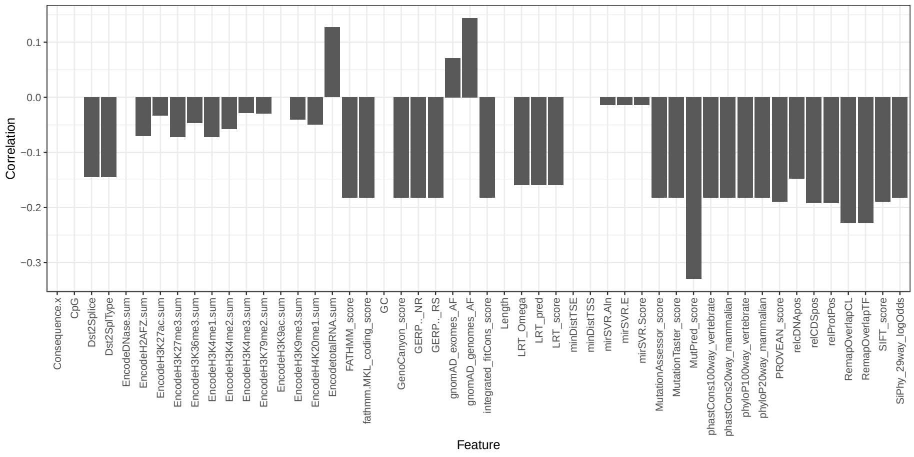


Figure 8: Correlations of features' missingness indicators to positive outcome indicator.

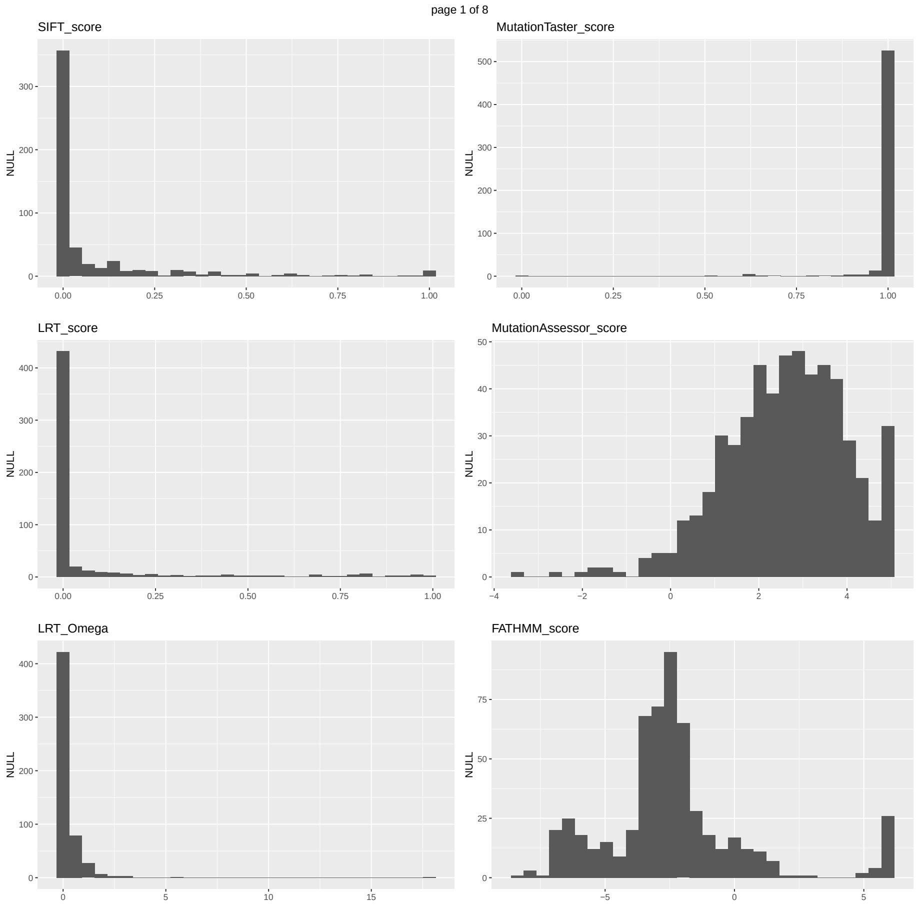


Figure 9: Histograms of the observed values of features, page 1.

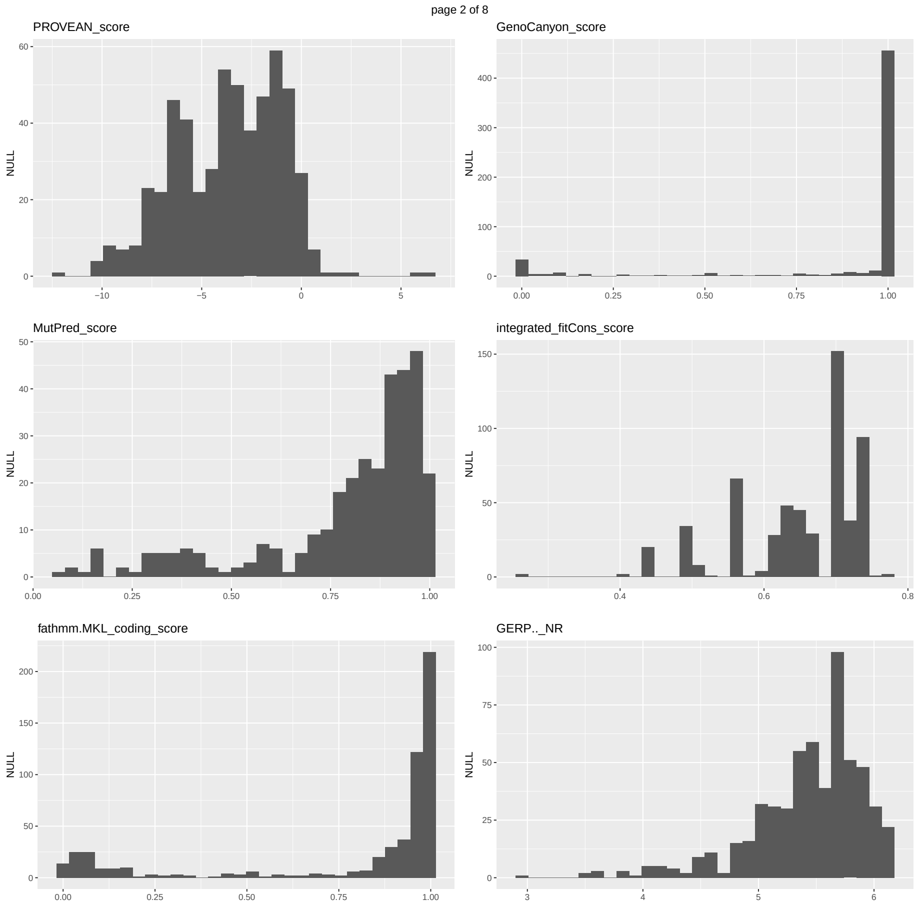


Figure 10: Histograms of the observed values of features, page 2.

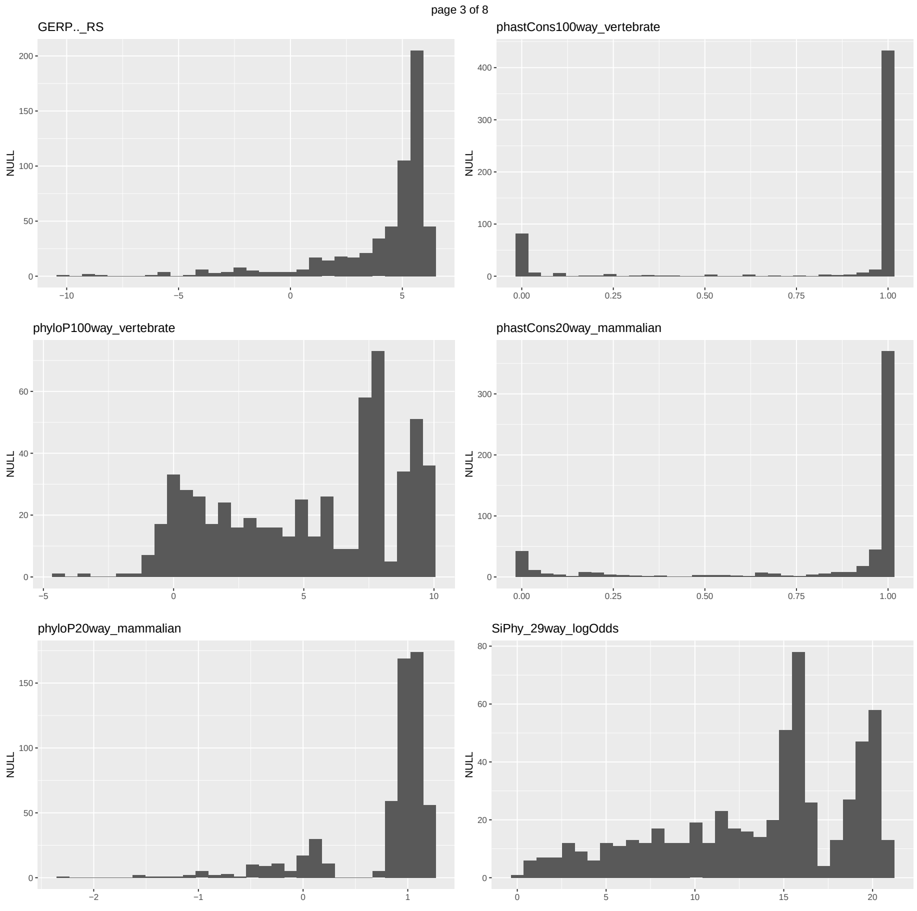


Figure 11: Histograms of the observed values of features, page 3.

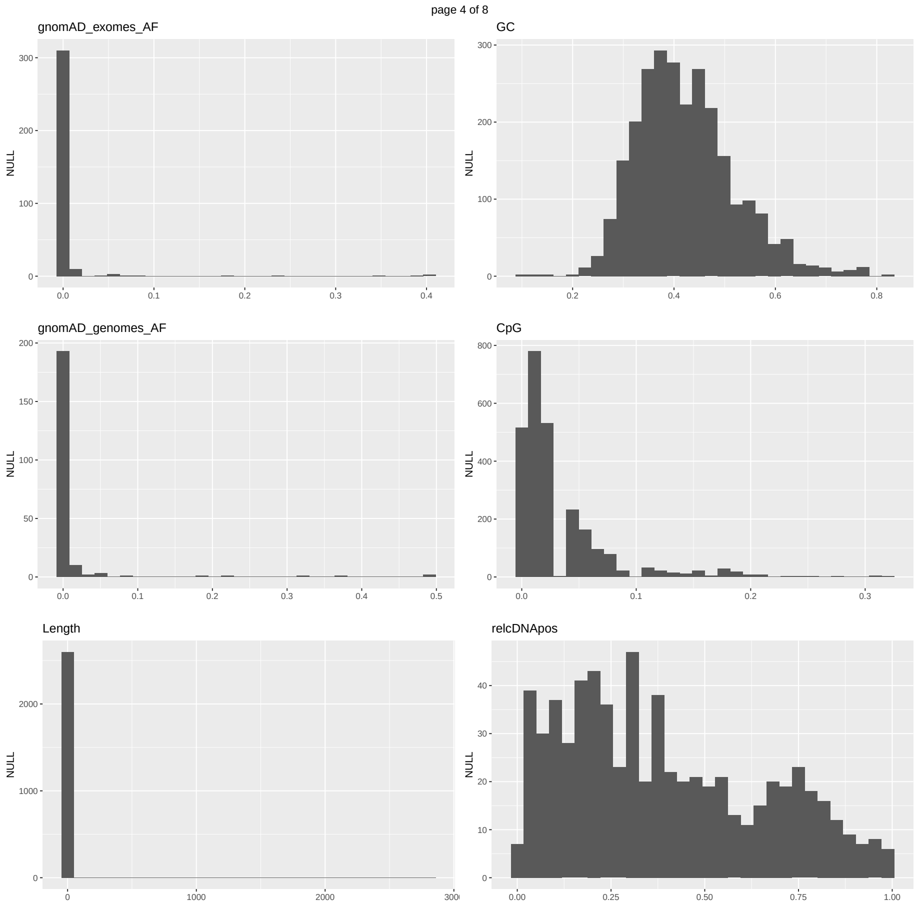


Figure 12: Histograms of the observed values of features, page 4.

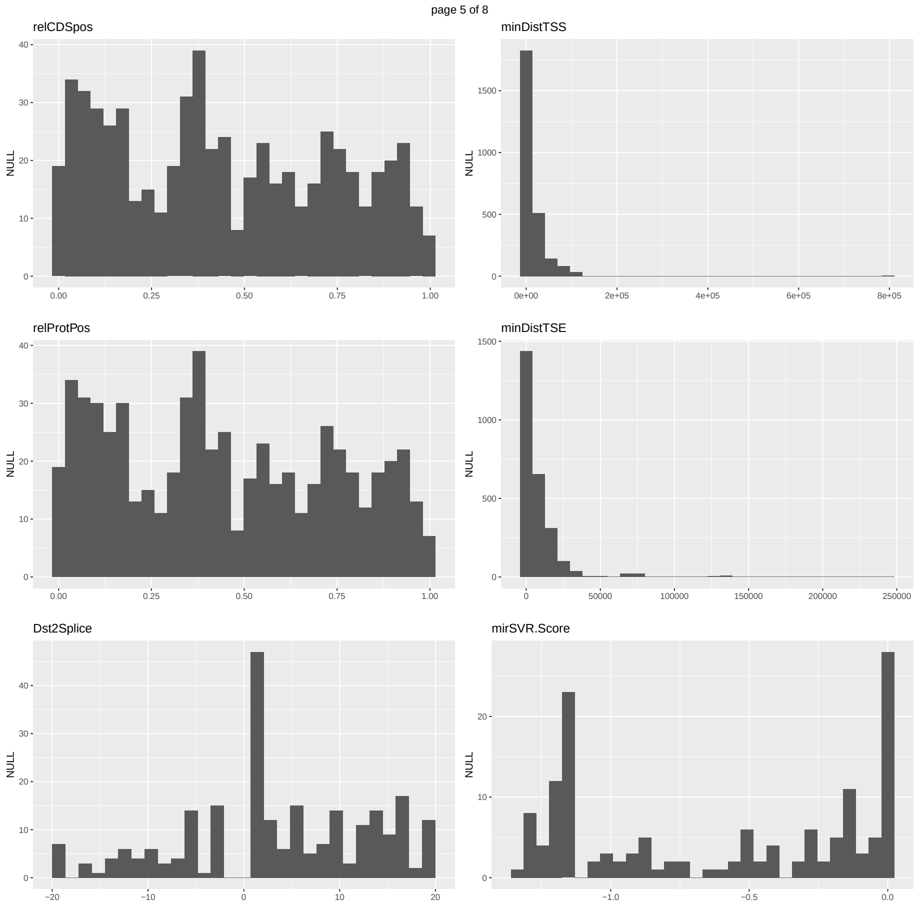


Figure 13: Histograms of the observed values of features, page 5.

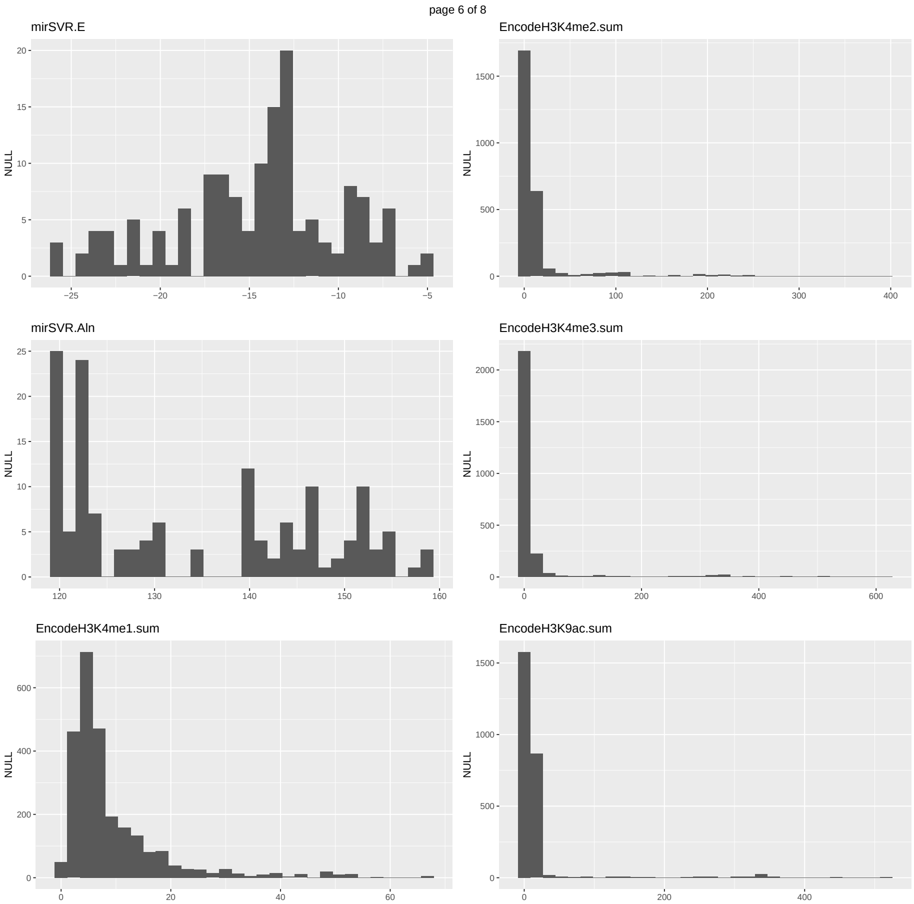


Figure 14: Histograms of the observed values of features, page 6.

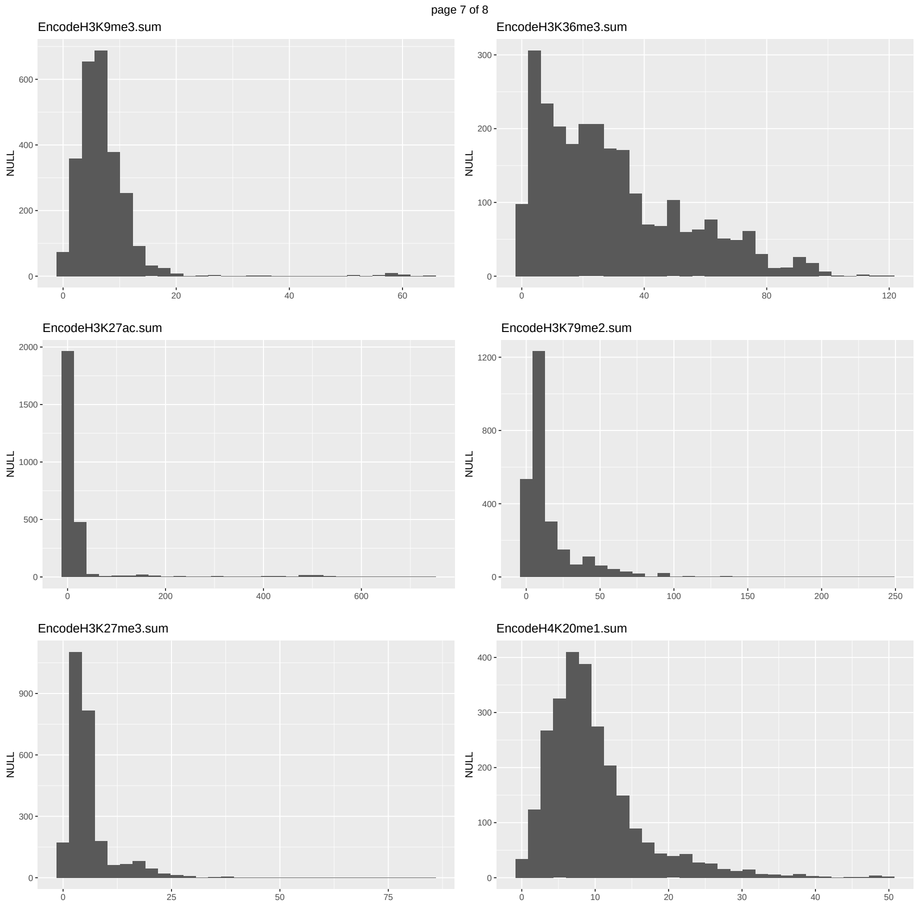


Figure 15: Histograms of the observed values of features, page 7.

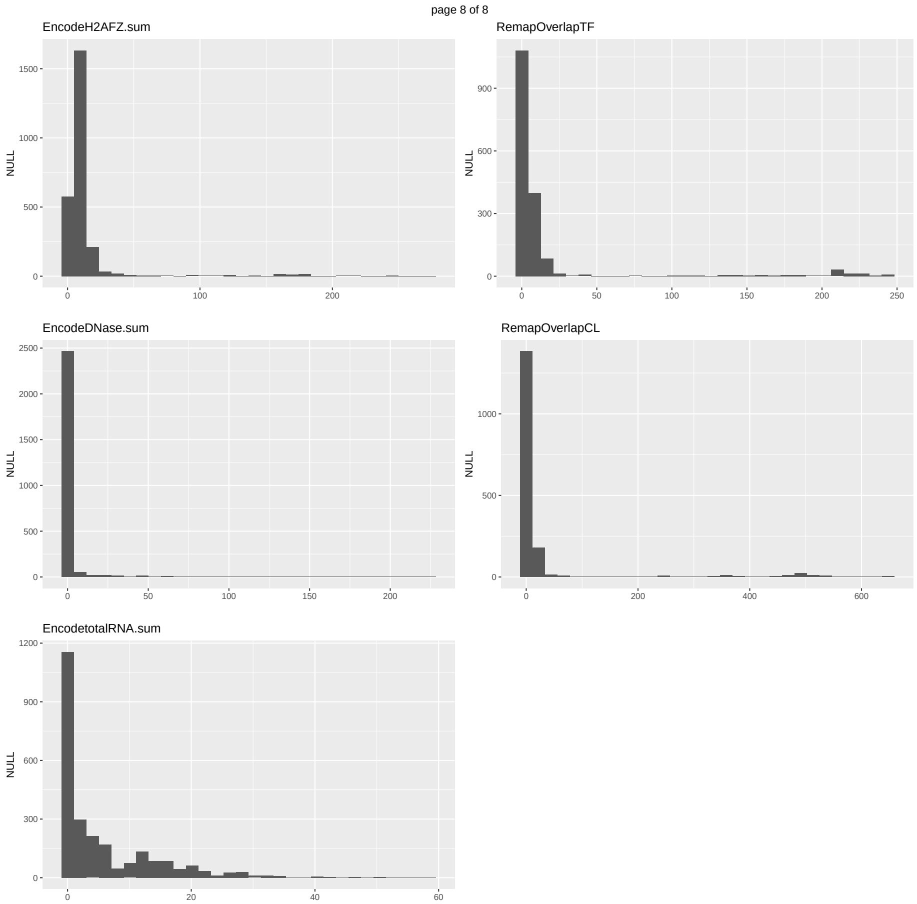


Figure 16: Histograms of the observed values of features, page 8.
